# Robust T cell immunity in convalescent individuals with asymptomatic or mild COVID-19

**DOI:** 10.1101/2020.06.29.174888

**Authors:** Takuya Sekine, André Perez-Potti, Olga Rivera-Ballesteros, Kristoffer Strålin, Jean-Baptiste Gorin, Annika Olsson, Sian Llewellyn-Lacey, Habiba Kamal, Gordana Bogdanovic, Sandra Muschiol, David J. Wullimann, Tobias Kammann, Johanna Emgård, Tiphaine Parrot, Elin Folkesson, Olav Rooyackers, Lars I. Eriksson, Anders Sönnerborg, Tobias Allander, Jan Albert, Morten Nielsen, Jonas Klingström, Sara Gredmark-Russ, Niklas K. Björkström, Johan K. Sandberg, David A. Price, Hans-Gustaf Ljunggren, Soo Aleman, Marcus Buggert, Karolinska COVID-19 Study Group

**Affiliations:** Center for Infectious Medicine, Department of Medicine Huddinge, Karolinska Institutet, Karolinska University Hospital, Stockholm, Sweden; Division of Infectious Diseases, Karolinska University Hospital and Department of Medicine Huddinge, Karolinska Institutet, Stockholm, Sweden; Division of Infection and Immunity, Cardiff University School of Medicine, University Hospital of Wales, Cardiff, UK; Department of Clinical Microbiology, Karolinska University Hospital, Stockholm, Sweden; Department of Clinical Interventions and Technology, Karolinska Institutet, Stockholm, Sweden; Function Perioperative Medicine and Intensive Care, Karolinska University Hospital, Stockholm, Sweden; Department of Physiology and Pharmacology, Karolinska Institutet, Stockholm, Sweden; Division of Clinical Microbiology, Department of Laboratory Medicine, Karolinska Institute, Karolinska University Hospital, Stockholm, Sweden; Department of Microbiology, Tumor and Cell Biology, Karolinska Institutet, Stockholm, Sweden; Department of Health Technology, Technical University of Denmark, DK-2800 Lyngby, Denmark; Instituto de Investigaciones Biotecnológicas, Universidad Nacional de San Martín, San Martín, Argentina; Systems Immunity Research Institute, Cardiff University School of Medicine, University Hospital of Wales, Cardiff, UK

## Abstract

SARS-CoV-2-specific memory T cells will likely prove critical for long-term immune protection against COVID-19. We systematically mapped the functional and phenotypic landscape of SARS-CoV-2-specific T cell responses in a large cohort of unexposed individuals as well as exposed family members and individuals with acute or convalescent COVID-19. Acute phase SARS-CoV-2-specific T cells displayed a highly activated cytotoxic phenotype that correlated with various clinical markers of disease severity, whereas convalescent phase SARS-CoV-2-specific T cells were polyfunctional and displayed a stem-like memory phenotype. Importantly, SARS-CoV-2-specific T cells were detectable in antibody-seronegative family members and individuals with a history of asymptomatic or mild COVID-19. Our collective dataset shows that SARS-CoV-2 elicits robust memory T cell responses akin to those observed in the context of successful vaccines, suggesting that natural exposure or infection may prevent recurrent episodes of severe COVID-19 also in seronegative individuals.

## INTRODUCTION

The world changed in December 2019 with the emergence of a new zoonotic pathogen, severe acute respiratory syndrome coronavirus 2 (SARS-CoV-2), which causes a variety of clinical syndromes collectively termed coronavirus disease 2019 (COVID-19). At present, there is no vaccine against SARS-CoV-2, and the excessive inflammation associated with severe COVID-19 can lead to respiratory failure, septic shock, and ultimately, death (Guan et al., 2020; Wolfel et al., 2020; Wu and McGoogan, 2020). The overall mortality rate is 0.5–3.5% (Guan et al., 2020; Wolfel et al., 2020; Wu and McGoogan, 2020). However, most people seem to be affected less severely and either remain asymptomatic or develop only mild symptoms during COVID-19 (He et al., 2020b; Wei et al., 2020; Yang et al., 2020). It will therefore be critical in light of the ongoing pandemic to determine if people with milder forms of COVID-19 develop robust immunity against SARS-CoV-2.

Global efforts are currently underway to map the determinants of immune protection against SARS-CoV-2. Recent data have shown that SARS-CoV-2 infection generates near-complete protection against rechallenge in rhesus macaques (Chandrashekar et al., 2020), and similarly, there is limited evidence of reinfection in humans with previously documented COVID-19 (Kirkcaldy et al., 2020). Further work is therefore required to define the mechanisms that underlie these observations and evaluate the durability of protective immune responses elicited by primary infection with SARS-CoV-2. Most correlative studies of immune protection against SARS-CoV-2 have focused on the induction of neutralizing antibodies (Hotez et al., 2020; Robbiani et al., 2020; Seydoux et al., 2020; Wang et al., 2020). However, antibody responses are not detectable in all patients, especially those with less severe forms of COVID-19 (Long et al., 2020; Mallapaty, 2020; Woloshin et al., 2020). Previous work has also shown that memory B cell responses tend to be short-lived after infection with SARS-CoV-1 (Channappanavar et al., 2014; Tang et al., 2011). In contrast, memory T cell responses can persist for many years (Nina Le Bert, 2020; Tang et al., 2011; Yang et al., 2006) and, in mice, protect against lethal challenge with SARS-CoV-1 (Channappanavar et al., 2014).

SARS-CoV-2-specific T cells have been identified in humans (Grifoni et al., 2020; Ni et al., 2020). It has nonetheless remained unclear to what extent various features of the T cell immune response associate with antibody responses and the clinical course of acute and convalescent COVID-19. To address this knowledge gap, we characterized SARS-CoV-2-specific CD4^+^ and CD8^+^ T cells in outcome-defined cohorts of donors (total n = 203) from Sweden, which has used a more “open” strategy, and as such durable spread, of COVID-19 than many other countries in Europe (Habib, 2020).

## RESULTS

Our preliminary analyses showed that the absolute numbers and relative frequencies of CD4^+^ and CD8^+^ T cells were unphysiologically low in patients with acute moderate or severe COVID-19 (Figure 1A and Figure S2A, B). This finding has been reported previously (He et al., 2020a; Liu et al., 2020). We then used a 31-parameter flow cytometry panel to assess the phenotypic landscape of these immune perturbations in direct comparisons with healthy blood donors and individuals who had recovered from asymptomatic/mild COVID-19 acquired early during the pandemic (February to March 2020). Unbiased principal component analysis (PCA) revealed a clear segregation between memory T cells from patients with acute moderate or severe COVID-19 and memory T cells from convalescent individuals and healthy blood donors (Figure 1B), driven largely by the expression of CD38, CD69, Ki-67, and programmed cell death protein 1 (PD-1) in the CD4^+^ compartment and by the expression of CD38, CD39, CD69, cytotoxic T-lymphocyte-associated protein 4 (CTLA-4), human leukcoyte antigen (HLA)-DR, Ki-67, lymphocyte-activation gene 3 (LAG-3), and T cell immunoglobulin and mucin domain-containing protein 3 (TIM-3) in the CD8^+^ compartment (Figure 1B, C and Figure S2C).

**Figure 1.**
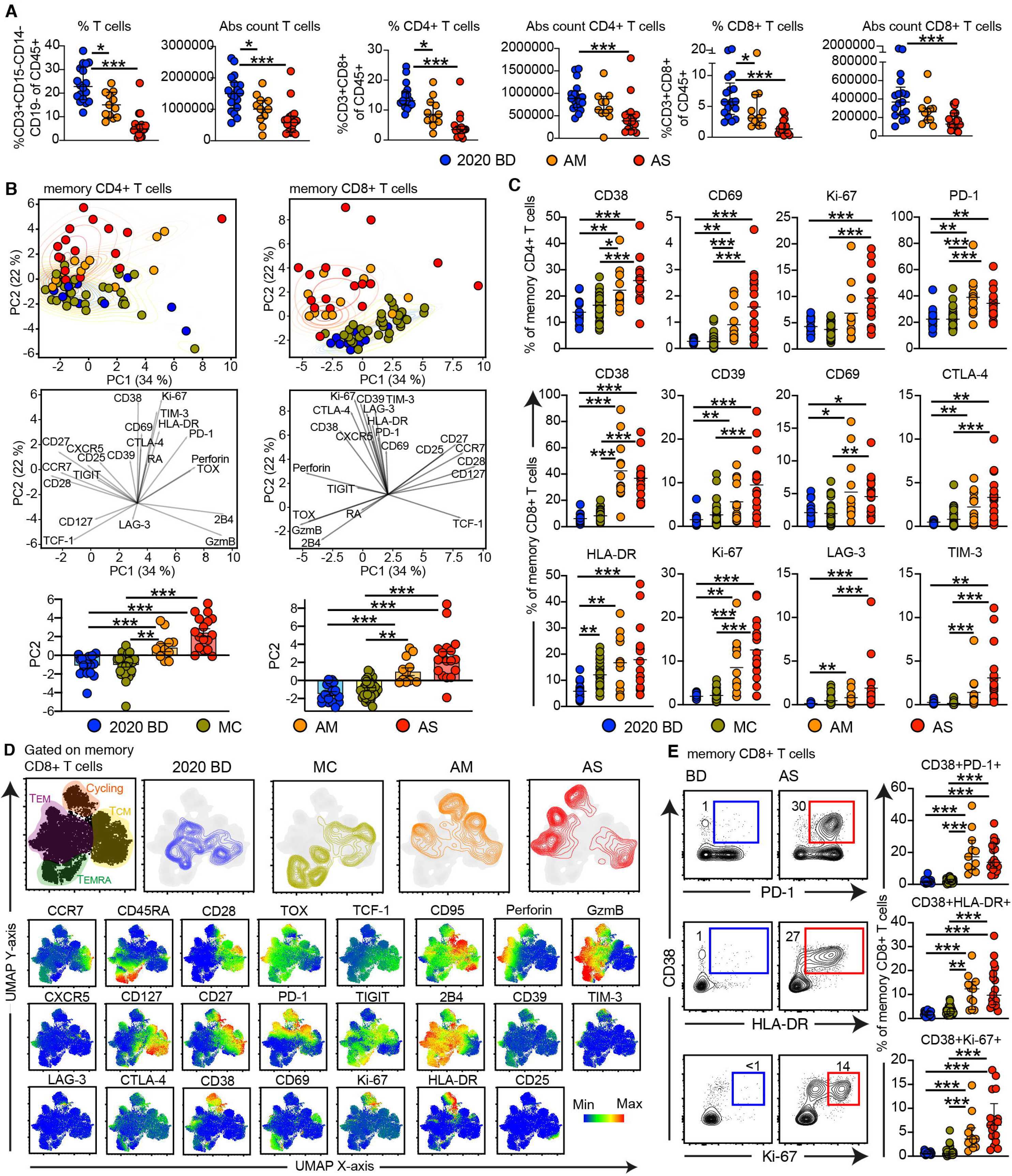
T cell perturbations in COVID-19. (**A**) Dot plots summarizing the absolute counts and relative frequencies of CD3^+^ (left), CD4^+^ (middle), and CD8^+^ T cells (right) in healthy blood donors from 2020 (2020 BD) and patients with acute moderate (AM) or severe COVID-19 (AS). Each dot represents one donor. Data are shown as median ± IQR. (**B**) Top: PCA plot showing the distribution and segregation of memory CD4^+^ and CD8^+^ T cells by group. MC: individuals in the convalescent phase after asymptomatic/mild COVID-19. Each dot represents one donor. Memory cells were defined by exclusion of naive cells (CCR7^+^ CD45RA^+^ CD95^−^). Middle: PCA plots showing the corresponding trajectories of key markers that influenced the group-defined segregation of memory CD4^+^ and CD8^+^ T cells. Bottom: dot plot showing the group-defined distribution of markers in PC2. Each dot represents one donor. (**C**) Dot plots summarizing the frequencies of CD4^+^ (top) and CD8^+^ T cells (bottom) expressing the indicated activation/cycling markers. Each dot represents one donor. Data are shown as median ± IQR. (**D**) Top: UMAP plots showing the clustering of memory CD8^+^ T cells by group in relation to all memory CD8^+^ T cells (left). Bottom: UMAP plots showing the expression of individual markers (n = 3 donors per group). (**E**) Left: representative flow cytometry plots showing the expression of activation/cycling markers among CD8^+^ T cells by group. Numbers indicate percentages in the drawn gates. Right: dot plots showing the expression frequencies of activation/cycling markers among memory CD8^+^ T cells by group. Key as in B. Each dot represents one donor. Data are shown as median ± IQR. \**P* < 0.05, \*\**P* < 0.01, \*\*\**P* < 0.001.

To extend these findings, we concatenated all memory CD4^+^ T cells (Figure S3A) and memory CD8^+^ T cells (Figure 1D) from healthy blood donors, convalescent individuals, and patients with acute moderate or severe COVID-19 via Uniform Manifold Approximation and Projection (UMAP). Distinct topographical clusters were apparent in each group (Figure 1D and S3A). In particular, memory CD4^+^ T cells (Figure S3A) and memory CD8^+^ T cells (Figure 1D) from patients with acute moderate or severe COVID-19 expressed a distinct cluster of markers associated with activation and the cell cycle, including CD38, HLA-DR, Ki-67, and PD-1. This finding was confirmed via manual gating of the flow cytometry data (Figure 1E). Correlative analyses further demonstrated that the activated/cycling phenotype was strongly associated with various clinical parameters, including age, hemoglobin concentration, platelet count, and plasma levels of alanine aminotransferase, albumin, D-dimer, fibrinogen, and myoglobin (Figure S3B and S3C), but less strongly associated with plasma levels of various inflammatory markers (Figure S4).

In most donors with acute COVID-19, we observed a pattern of increased CD38 expression, also without HLA-DR, Ki-67 and PD-1 expression (Figure S5A and S5B), compared to healthy blood donors. We confirmed that CD8^+^ T cells specific for cytomegalovirus (CMV) or Epstein-Barr virus (EBV) expressed increased frequencies of CD38, indicating that single CD38 expression could be driven by inflammation or other features in COVID-19 (Figure 2A, B and Figure S5C). Notably though, CMV- and EBV-specific CD8^+^ T cells did not express elevated of HLA-DR, Ki-67, or PD-1 and/or in combination with CD38, during acute moderate or severe COVID-19 compared with convalescent individuals and healthy blood donors, indicating limited bystander proliferation and activation during the early phase of infection with SARS-CoV-2 (Figure 2A, B and Figure S5C). Actively proliferating CD8^+^ T cells, defined by the expression of Ki-67, instead exhibited a predominant CCR7^−^ CD27^+^ CD28^+^ CD45RA^−^ CD127^−^ phenotype in patients with acute moderate or severe COVID-19 (Figure S5D), as reported previously in the context of vaccination and other viral infections (Buggert et al., 2018b; Miller et al., 2008). On the basis of these findings, we used overlapping peptides spanning the immunogenic domains of the SARS-CoV-2 membrane, nucleocapsid, and spike proteins to stimulate peripheral blood mononuclear cells (PBMCs) from patients with acute moderate or severe COVID-19, and found that responding CD4^+^ and CD8^+^ T cells displayed an activated/cycling (CD38^+^ HLA-DR^+^ Ki67^+^ PD-1^+^) phenotype (Figure 2C). These results were confirmed using an activation-induced marker (AIM) assay to measure the upregulation of CD69 and 4-1BB (CD137), which suggests that most CD38^+^ PD-1^+^ CD8^+^ T cells were specific for SARS-CoV-2 (Figure 2D).

**Figure 2.**
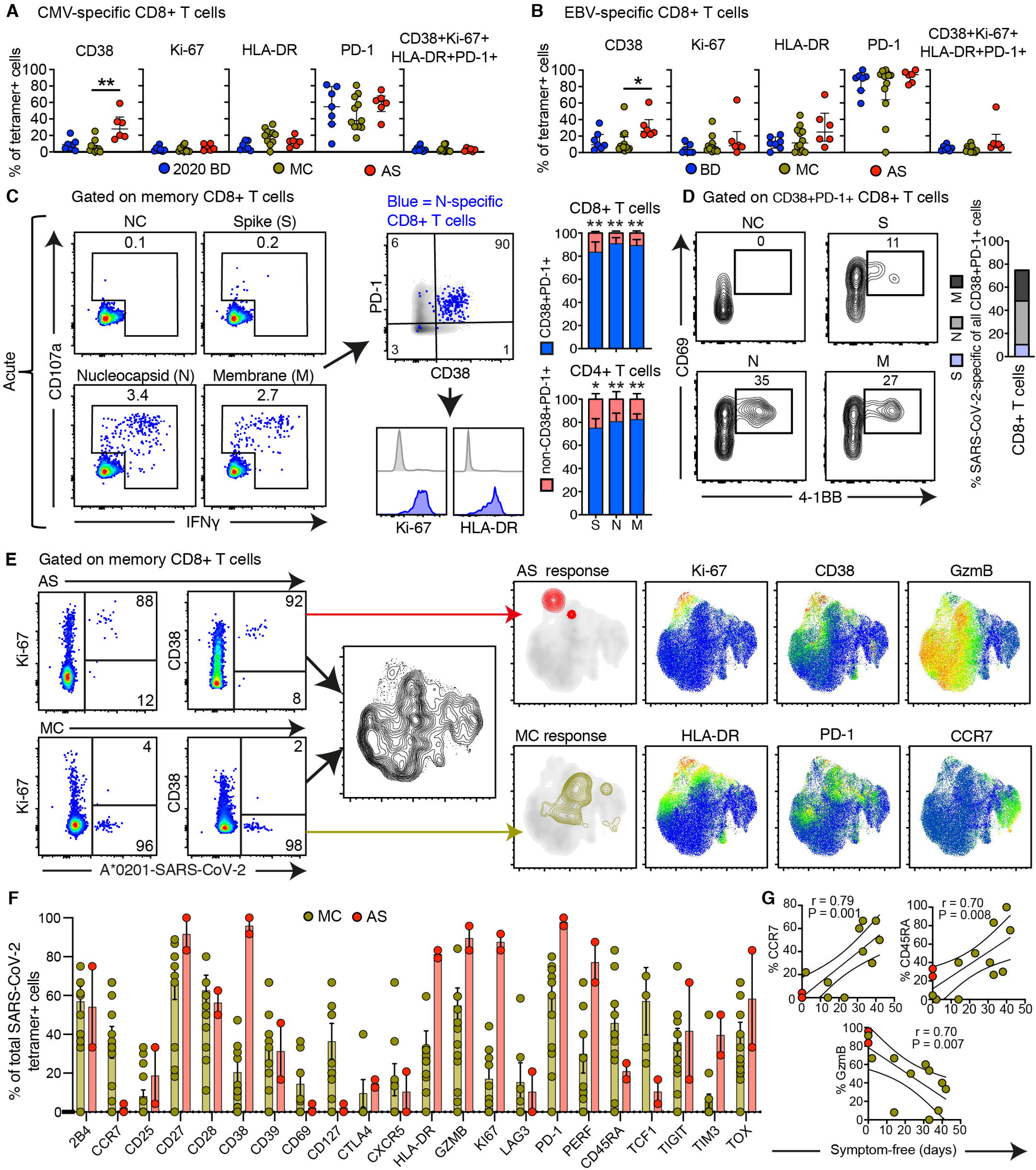
Phenotypic characteristics of SARS-CoV-2-specific T cells in acute and convalescent COVID-19. (**A** and **B**) Dot plots showing the expression frequencies of activation/cycling markers among tetramer^+^ CMV-specific (A) or EBV-specific CD8^+^ T cells (B) by group. Each dot represents one specificity in one donor. Data are shown as median ± IQR. (**C**) Representative flow cytometry plots (left) and bar graphs (right) showing the expression of activation/cycling markers among CD107a^+^ and/or IFN-γ^+^ SARS-CoV-2-specific CD4^+^ and CD8^+^ T cells (n = 6 donors). Numbers indicate percentages in the drawn gates. NC: negative control. (**D**) Representative flow cytometry plots (left) and bar graph (right) showing the upregulation of CD69 and 4-1BB (AIM assay) among CD38^+^ PD-1^+^ SARS-CoV-2-specific CD8^+^ T cells (n = 6 donors). Numbers indicate percentages in the drawn gates. (**E**) Left: representative flow cytometry plots showing the expression of activation/cycling markers among tetramer^+^ SARS-CoV-2-specific CD8^+^ T cells by group. Middle: UMAP plot showing the clustering of memory CD8^+^ T cells. Right: UMAP plots showing the clustering of tetramer^+^ SARS-CoV-2-specific CD8^+^ T cells by group and the expression of individual markers (n = 2 donors). (**F**) Bar graph showing the expression frequencies of all quantified markers among tetramer^+^ SARS-CoV-2-specific CD8^+^ T cells by group. Each dot represents combined specificities in one donor. Data are shown as median ± IQR. (**G**) Bivariate plots showing the pairwise correlations between symptom-free days and the expression frequencies of CCR7, CD45RA, or granzyme B (GzmB). Each dot represents combined specificities in one donor. Key as in F. \**P* < 0.05, \*\**P* < 0.01, \*\*\**P* < 0.001.

In further experiments, we used HLA class I tetramers as probes to detect CD8^+^ T cells specific for predicted optimal epitopes from SARS-CoV-2 (Table S2). A vast majority of tetramer^+^ CD8^+^ T cells in the acute phase of infection, but not during convalescence, displayed an activated/cycling phenotype (Figure 2E). In general, early SARS-CoV-2-specific CD8^+^ T cell populations were characterized by the expression of immune activation molecules (CD38, HLA-DR, Ki-67), inhibitory receptors (PD-1, TIM-3), and cytotoxic molecules (granzyme B, perforin), whereas convalescent phase SARS-CoV-2-specific CD8^+^ T cell populations were skewed toward an early differentiated memory (CCR7^+^ CD127^+^ CD45RA^+^ TCF-1^+^) phenotype (Figure 2F). Importantly, the expression frequencies of CCR7 and CD45RA among SARS-CoV-2-specific CD8^+^ T cells were positively correlated with the number of symptom-free days after infection, whereas the expression frequency of granzyme B among SARS-CoV-2-specific CD8^+^ T cells was inversely correlated with the number of symptom-free days after infection (Figure 2G). Time from exposure was therefore associated with the emergence of stem-like memory SARS-CoV-2-specific CD8^+^ T cells.

On the basis of these observations, we quantified functional SARS-CoV-2-specific memory T cell responses across five distinct cohorts, including healthy individuals who donated blood either before or during the pandemic, family members who shared a household with convalescent individuals and were exposed at the time of symptomatic disease, and individuals in the convalescent phase after asymptomatic/mild or severe COVID-19. We detected potentially cross-reactive T cell responses directed against the membrane and spike proteins in healthy individuals who donated blood before the pandemic, consistent with previous reports (Grifoni et al., 2020; Nina Le Bert, 2020), but nucleocapsid reactivity was notably absent in this cohort (Figure 3A and S6A, S6B). The highest response frequencies across all three proteins were observed in convalescent individuals who experienced severe COVID-19. Progressively lower response frequencies were observed in convalescent individuals with a history of asymptomatic/mild COVID-19, exposed family members, and healthy individuals who donated blood during the pandemic (Figure 3A).

**Figure 3.**
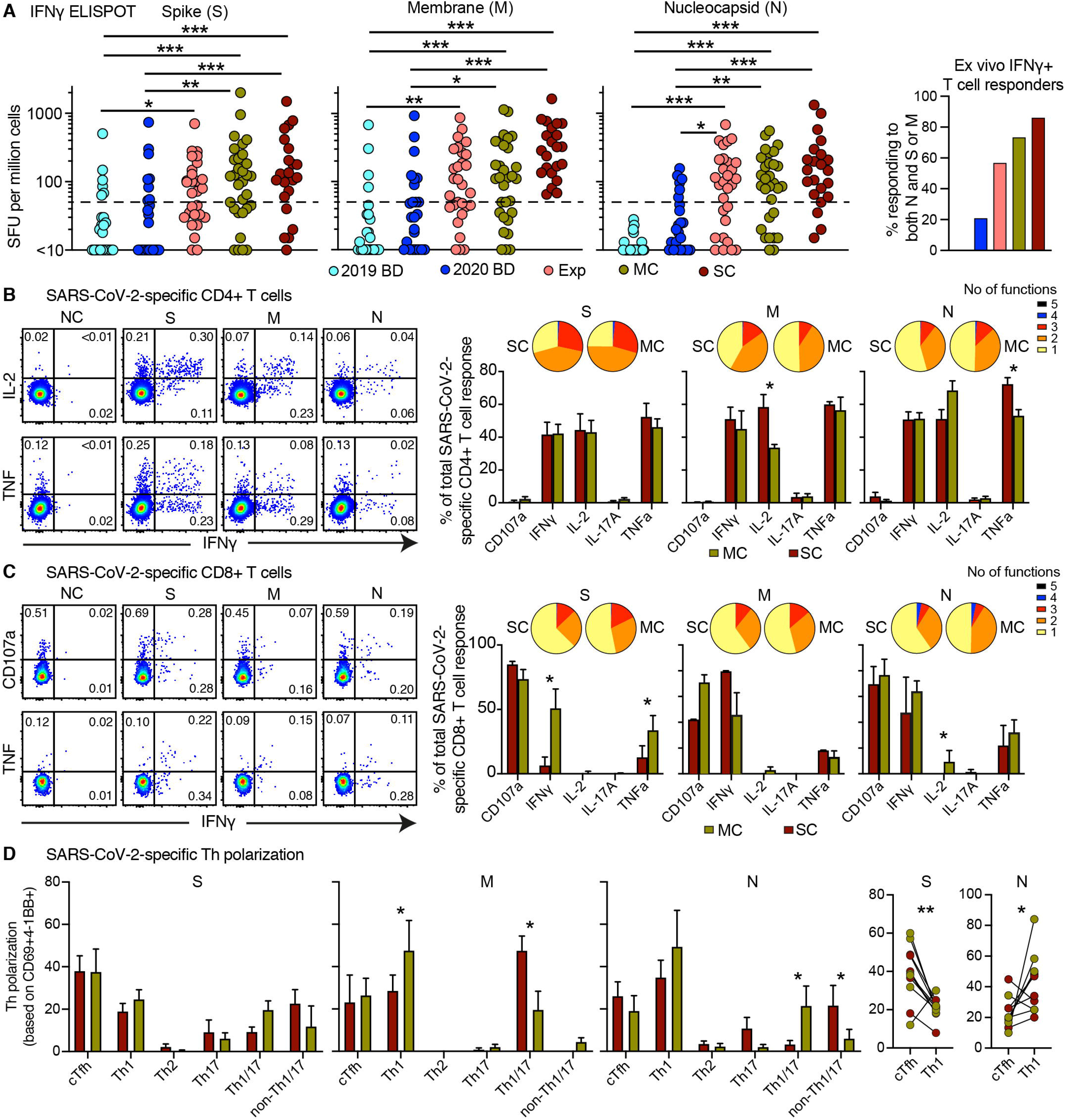
Functional characteristics of SARS-CoV-2-specific T cells in convalescent COVID-19. (**A**) Left: dot plots showing the frequencies of IFN-γ-producing cells responding to overlapping peptides spanning the immunogenic domains of the SARS-CoV-2 membrane (M), nucleocapsid (N), and spike proteins (S) by group (ELISpot assays). Each dot represents one donor. The dotted line indicates the cut-off for positive responses. Right: bar graph showing the frequencies of IFN-γ-producing cells responding to both the internal (N) and surface antigens (M and/or S) of SARS-CoV-2 by group (ELISpot assays). BD 2019: healthy blood donors from 2019. Exp: exposed family members. SC: individuals in the convalescent phase after severe or (MC) asymptomatic/mild COVID-19. SFU: spot-forming unit. (**B** and **C**) Left: representative flow cytometry plots showing the functional profiles of SARS-CoV-2-specific CD4^+^ (B) and CD8^+^ T cells (C) from a convalescent individual (group MC). Numbers indicate percentages in the drawn gates. Right: bar graphs showing the distribution of individual functions among SARS-CoV-2-specific CD4^+^ (B) and CD8^+^ T cells (C) from convalescent individuals in groups MC (n = 12) or SC (n = 14). Key as in A. Data are shown as median ± IQR. (**D**) Left: bar graphs showing the functional polarization of SARS-CoV-2-specific CD4^+^ T from convalescent individuals in groups MC (n = 8) and SC (n = 8). Subsets were defined as CXCR5^+^ (cTfh), CCR4^−^ CCR6^−^ CXCR3^+^ CXCR5^−^ (Th1), CCR4^+^ CCR6^−^ CXCR3^−^ CXCR5^−^ (Th2), CCR4^−^ CCR6^+^ CXCR3^−^ CXCR5^−^ (Th17), CCR4^−^ CCR6^+^ CXCR3^+^ CXCR5^−^ (Th1/17), and CCR4^−^ CCR6^−^ CXCR3^−^ CXCR5^−^ (non-Th1/2/17). Data are shown as median ± IQR. Right: line graph comparing cTfh *versus* Th1 polarization by specificity in convalescent individuals from groups MC and SC. Each dot represents one donor. \**P* < 0.05, \*\**P* < 0.01, \*\*\**P* < 0.001.

To assess the functional capabilities of SARS-CoV-2-specific memory CD4^+^ and CD8^+^ T cells in convalescent individuals, we stimulated PBMCs with the overlapping membrane, nucleocapsid, and spike peptide sets and measured a surrogate marker of degranulation (CD107a) along with the production of interferon (IFN)-γ, IL-2, and TNF (Figure 3B, C). SARS-CoV-2-specific CD4^+^ T cells predominantly expressed IFN-γ, IL-2, and TNF (Figure 3B), whereas SARS-CoV-2-specific CD8^+^ T cells predominantly expressed IFN-γ and TNF and mobilized CD107a (Figure 3C). We then used the AIM assay to determine the functional polarization of SARS-CoV-2-specific CD4^+^ T cells. Interestingly, spike-specific CD4^+^ T cells were skewed toward a cTfh profile, whereas membrane-specific and nucleocapsid-specific CD4^+^ T cells were skewed toward a Th1 or a Th1/Th17 profile (Figure 3D and S7A, S7B).

In the next set of experiments, we assessed the recall capabilities of SARS-CoV-2-specific CD4^+^ and CD8^+^ T cells in convalescent individuals, exposed family members, and healthy blood donors. Proliferative responses were identified by tracking the progressive dilution of a cytoplamsic dye (CellTrace Violet; CTV) after stimulation with the overlapping membrane, nucleocapsid, and spike peptide sets, and functional responses to the same antigens were evaluated 5 days later by measuring the production of IFN-γ (Blom et al., 2013; Buggert et al., 2014a). Anamnestic responses in the CD4^+^ and CD8^+^ T cell compartments, quantified as a function of CTV^low^ IFN-γ^+^ events (Figure 4A), were detected in most convalescent individuals and exposed family members (Figure 4B, C). SARS-CoV-2-specific CD4^+^ T cell responses were proportionately larger overall than the corresponding SARS-CoV-2-specific CD8^+^ T cell responses (Figure 4D). In addition, most IFN-γ^+^ SARS-CoV-2-specific CD4^+^ T cells produced TNF, and most IFN-γ^+^ SARS-CoV-2-specific CD8^+^ T cells produced granzyme B and perforin (Figure 4E).

**Figure 4.**
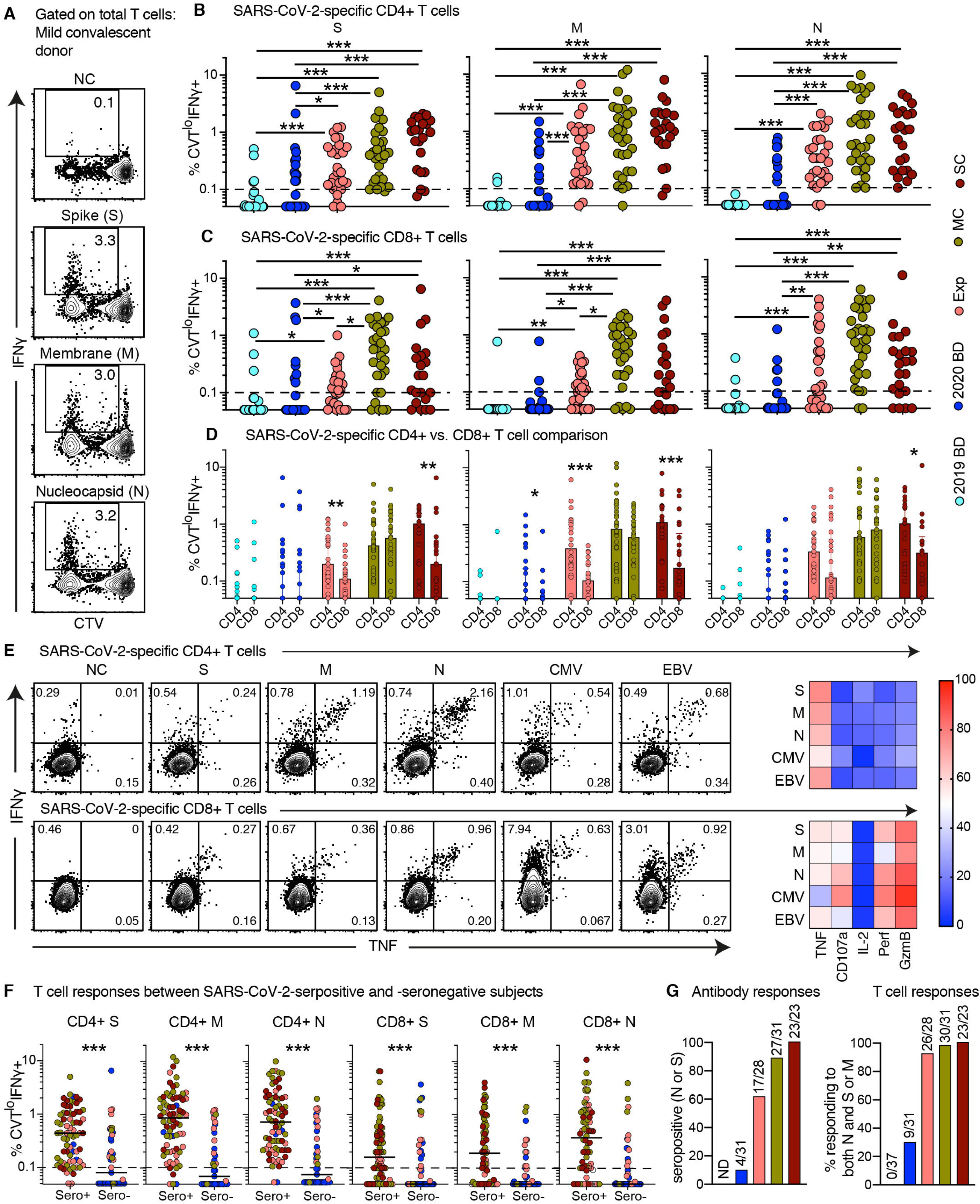
Proliferative capabilities of SARS-CoV-2-specific T cells in convalescent COVID-19. (**A**) Representative flow cytometry plots showing the proliferation (CTV^−^) and functionality (IFN-γ^+^) of SARS-CoV-2-specific T cells from a convalescent individual (group MC) after stimulation with overlapping peptides spanning the immunogenic domains of the SARS-CoV-2 membrane (M), nucleocapsid (N), and spike proteins (S). Numbers indicate percentages in the drawn gates. (**B** and **C**) Dot plots showing the frequencies of CTV^−^ IFN-γ^+^ SARS-CoV-2-specific CD4^+^ (B) and CD8^+^ T cells (C) by group and specificity. Each dot represents one donor. The dotted line indicates the cut-off for positive responses. (**D**) Bar graphs comparing the frequencies of CTV^−^ IFN-γ^+^ SARS-CoV-2-specific CD4^+^ *versus* CD8^+^ T cells by group and specificity. Each dot represents one donor. Data are shown as median ± IQR. (**E**) Left: representative flow cytometry plots showing the production of IFN-γ and TNF among CTV^−^ virus-specific CD4^+^ (top) and CD8^+^ T cells (bottom) from a convalescent individual (group MC). Numbers indicate percentages in the drawn gates. Right: heatmaps summarizing the functional profiles of CTV^−^ IFN-γ^+^ virus-specific CD4^+^ (top) and CD8^+^ T cells (bottom). Data are shown as mean frequencies (key). (**F**) Dot plots showing the frequencies of CTV^−^ IFN-γ^+^ SARS-CoV-2-specific CD4^+^ and CD8^+^ T cells by group, serostatus, and specificity. Each dot represents one donor. The dotted line indicates the cut-off for positive responses. Key as in B. **(G)** Left: bar graph showing percent seropositivity by group. Right: bar graph showing the percentage of individuals in each group with detectable T cell responses directed against both the internal (N) and surface antigens (M and/or S) of SARS-CoV-2. *P < 0.05, **P < 0.01, ***P < 0.001.

In a final set of analyses, we compared the SARS-CoV-2-specific antibody and T cell responses in and between the different groups. The anti-SARS-CoV-2 IgG responses against the nucleocapsid and spike antigens were strongly correlated (Figure S8A). Further analysis revealed that SARS-CoV-2-specific CD4^+^ and CD8^+^ T cell responses were present in seronegative individuals, albeit at lower frequencies compared with seropositive individuals (Figure 4F). These discordant responses were nonetheless pronounced in some convalescent individuals with a history of asymptomatic/mild COVID-19, exposed family members, and healthy individuals who donated blood during the pandemic (Figure 4F and S8B, S8C), often targeting both the internal (nucleocapsid) and surface antigens (membrane and/or spike) of SARS-CoV-2 (Figure 4G). Potent memory T cell responses were therefore elicited in the absence or presence of circulating antibodies, consistent with a non-redundant role as key determinants of immune protection against COVID-19 (Chandrashekar et al., 2020).

## DISCUSSION

We are currently facing the biggest global health emergency in decades, namely the devastating outbreak of COVID-19. In the absence of a protective vaccine, it will be critical to determine if exposed and/or infected people, especially those with asymptomatic or very mild forms of the disease who likely act inadvertently as the major transmitters, develop robust adaptive immunity against SARS-CoV-2 (Long et al., 2020).

In this study, we used a systematic approach to map cellular and humoral immune responses against SARS-CoV-2 in patients with acute moderate or severe COVID-19, individuals in the convalescent phase after asymptomatic/mild or severe COVID-19, exposed family members, and healthy individuals who donated blood before (2019) or during the pandemic (2020). Individuals in the convalescent phase after asymptomatic/mild COVID-19 were traced after returning to Sweden from endemic areas (mostly Northern Italy). These donors exhibited robust memory T cell responses months after infection, even in the absence of detectable circulating antibodies specific for SARS-CoV-2, indicating a previously unanticipated degree of population-level immunity against COVID-19.

We found that T cell activation, characterized by the expression of CD38, was a hallmark of acute COVID-19. Similar findings have been reported previously in the absence of specificity data (Huang et al., 2020; Thevarajan et al., 2020; Wilk et al., 2020). Many of these T cells also expressed HLA-DR, Ki-67, and PD-1, indicating a combined activation/cycling phenotype, and expression levels of CD38 in particular correlated with disease severity, but notably not to a high degree to inflammatory markers. Our data also showed that many activated/cycling T cells in the acute phase were functionally replete and specific for SARS-CoV-2. Equivalent functional profiles have been observed early after immunization with successful vaccines (Blom et al., 2013; Miller et al., 2008; Precopio et al., 2007). Accordingly, the expression of multiple inhibitory receptors, including PD-1, likely indicates early activation rather than exhaustion (Zheng et al., 2020a; Zheng et al., 2020b).

Virus-specific memory T cells have been shown to persist for many years after infection with SARS-CoV-1 (Nina Le Bert, 2020; Tang et al., 2011; Yang et al., 2006). In line with these observations, we found that SARS-CoV-2-specific T cells acquired an early differentiated memory (CCR7^+^ CD127^+^ CD45RA^−/+^ TCF-1^+^) phenotype in the convalescent phase, as reported previously in the context of other viral infections and successful vaccines (Blom et al., 2013; Demkowicz et al., 1996; Fuertes Marraco et al., 2015; Precopio et al., 2007). This phenotype has been associated with stem-like properties (Betts et al., 2006; Blom et al., 2013; Demkowicz et al., 1996; Fuertes Marraco et al., 2015; Precopio et al., 2007). Accordingly, we found that SARS-CoV-2-specific T cells generated anamnestic responses to cognate antigens in the convalescent phase, characterized by extensive proliferation and polyfunctionality. Of particular note, we detected similar memory T cell responses directed against the internal (nucleocapsid) and surface proteins (membrane and/or spike) in some individuals lacking detectable circulating antibodies specific for SARS-CoV-2. Indeed, almost twice as many exposed family members and healthy individuals who donated blood during the pandemic generated memory T cell responses versus antibody responses, implying that seroprevalence as an indicator has underestimated the extent of population-level immunity against SARS-CoV-2.

It remains to be determined if a robust memory T cell response in the absence of detectable circulating antibodies can protect against SARS-CoV-2. This scenario has nonetheless been inferred from previous studies of MERS and SARS-CoV-1 (Channappanavar et al., 2014; Li et al., 2008; Zhao et al., 2017; Zhao et al., 2016), both of which have been shown to induce potent memory T cell responses that persist while antibody responses wane (Alshukairi et al., 2016; Shin et al., 2019; Tang et al., 2011). Moreover, vaccine-induced T cell responses, even in the absence of detectable antibodies, can protect mice against lethal challenge with SARS-CoV-1 (Channappanavar et al., 2014). In line with these observations, none of the convalescent individuals in this study, including those with previous asymptomatic/mild disease, have experienced further episodes of COVID-19.

Collectively, our data have provided a functional and phenotypic map of SARS-CoV-2-specific T cell immunity across the full spectrum of exposure, infection, and disease. The observation that most individuals with asymptomatic or mild COVID-19 generated highly functional durable memory T cell responses, not uncommonly in the relative absence of corresponding humoral responses, further suggested that natural exposure or infection could prevent recurrent episodes of severe COVID-19.

## MATERIALS AND METHODS

### Samples

Donors were assigned to one of seven groups for the purposes of this study. An eight-category NIH ordinal scale (defined below) and Sequential Organ Failure Assessment (SOFA) score were used to assess the severity of the disease at the highest point (Beigel et al., 2020; Singer et al., 2016). The NIH ordinal scale scores are as follows: 1, not hospitalized, no limitations of activities; 2, not hospitalized, limitation of activities, home oxygen requirement, or both; 3, hospitalized, not requiring supplemental oxygen and no longer requiring ongoing medical care (used if hospitalization was extended for infection-control reasons); 4, hospitalized, not requiring supplemental oxygen but requiring ongoing medical care (Covid-19–related or other medical conditions); 5, hospitalized, requiring any supplemental oxygen; 6, hospitalized, requiring noninvasive ventilation or use of high-flow oxygen devices; 7, hospitalized, receiving invasive mechanical ventilation or extracorporeal membrane oxygenation (ECMO); and 8, death.

Acute moderate (AM): patients requiring hospitalization and low-flow oxygen support of 0-3 L/min at sampling (n = 10). These patients had a median score of 5 (IQR 5-5) at NIH ordinal scale and a median Sequential Organ Failure Assessment (SOFA) score of 1 (IQR 1-1) at sampling. Acute severe (AS): patients requiring hospitalization in the high dependency unit or intensive care unit, with either low-flow oxygen support of > 10 L/min, high-flow oxygen support, or invasive mechanical ventilation (n = 17). 14 patients (82%) had required ECMO or invasive mechanical ventilation at intensive unit. These patients had a median NIH ordinal scale score of 7 (IQR 6-7) and a median SOFA score of 6 (IQR 3-6) at sampling. Mild Convalescent (MC): individuals in the convalescent phase after mild disease (n = 40). The majority were not hospitalized with mild symptoms (78%, n=31), while hospitalized patients had mild-moderate symptoms requiring either no oxygen support (n=7) or low intermittent oxygen support up to 1 L/min (n=2). These patients had a median of 1 (IQR 1-1) on the NIH ordinal scale. Severe Convalescent (SC): individuals in the convalescent phase after severe disease (n = 26). The median score was 6 (IQR 5-7) on NIH ordinal scale. Exposed (Exp): family members who shared a household with donors in groups MC or SC and were exposed at the time of symptomatic disease (n = 30), but without any diagnoses of COVID-19. These had all NIH ordinal scale of 1. 2020 Blood donors (BD): individuals who donated blood at the Karolinska University Hospital in May 2020 (during the pandemic; n = 55). 2019 BD: individuals who donated blood at the Karolinska University Hospital between July and September 2019 (before the pandemic; n = 25).

Individuals with acute COVID-19 were sampled 5–24 (median 14; IQR 11-17) days after the onset of symptoms debut and 1-8 (median 5; IQR 3-7) days after hospital admission (Table S1). All individuals with acute or convalescent disease tested positive for SARS-CoV-2 RNA. Group MC comprised individuals who had returned from endemic countries in Europe (mostly Northern Italy) between February and March 2020 and were among the first cases reported in Sweden. Seven persons in group Exp were negative for SARS-CoV-2 at the time of positive test for MC or SC donors, while rest were not tested. Individuals in groups 2020 BD, and 2019 BD were not tested for SARS-CoV-2 RNA.

All participants enrolled in this study provided written informed consent in accordance with protocols approved by the regional ethical research boards and the Declaration of Helsinki. Donor groups and clinical parameters are summarized in Table S1.

### Flow cytometry

PBMCs were isolated from venous blood samples via standard density gradient centrifugation and used immediately (groups AM, AS, MC, SC, Exp, and BD 2020) or after cryopreservation in liquid nitrogen (group BD 2019). Cells were washed in phosphate-buffered saline (PBS) supplemented with 2% fetal bovine serum (FBS) and 2 μM EDTA (FACS buffer) and stained with HLA class I tetramers and/or a directly conjugated antibody specific for CCR7 (clone G043H7; BioLegend) for 10 minutes at 37°C. Other surface markers were detected via the subsequent addition of directly conjugated antibodies at pretitrated concentrations for 20 minutes at room temperature, and viable cells were identified by exclusion using a LIVE/DEAD Fixable Aqua Dead Cell Stain Kit (Thermo Fisher Scientific). Cells were then washed again in FACS buffer and fixed/permeabilized using a FoxP3 / Transcription Factor Staining Buffer Set (eBioscience). Intracellular markers were detected via the addition of directly conjugated antibodies at pretitrated concentrations for 1 hour at 4°C. Stained cells were fixed in PBS containing 1% paraformaldehyde (Biotium) and stored at 4°C. Samples were acquired using a FACSymphony A5 (BD Biosciences). Data were analyzed with FlowJo software version 10.6.1 (FlowJo LLC). Gating strategies were based on fluorescent-minus-one or negative controls as described previously (Buggert et al., 2018a; Buggert et al., 2018b; Buggert et al., 2014b).

### Antibodies

Directly conjugated monoclonal antibodies with the following specificities were used in flow cytometry experiments: CCR4–PE (clone 1G1), CCR6–PE-Cy7 (clone 11A9), CD3–BUV805 (clones RPA-T8 or UCHT1), CD8–BUV395 (clone RPA-T8), CD25–PE-Cy5 (clone M-A251), CD28–BUV563 (clone CD28.2), CD38–BUV496 (clone HIT29), CD69–BV750 (clone FN50), CD95–BB630 (clone DX2), CD107a–PE-CF594 (clone H4A3), CTLA-4–BB755 (clone BNI3), CXCR5–APC-R700 (clone RF8B2), granzyme B–BB790 (clone GB11), HLA-DR–BUV615 (clone G46-6), IL-2–APC-R700 (clone MQ1-17H12), Ki-67–BB660 (clone B56), LAG-3–BUV661 (clone T47-530), perforin– BB700 (clone dG9), TIGIT–BUV737 (clone 741182), and 2B4–PE/Dazzle 594 (clone C1.7) from BD Biosciences; CCR7–APC-Cy7 (clone G043H7), CD14–BV510 (clone M5E2), CD19–BV510 (clone HIB19), CD27–BV785 (clone O323), CD39–BV711 (clone A1), CD45RA–BV421 or CD45RA–BV570 (clone HI100), CD127–BV605 (clone A019D5), CXCR3–AF647 (clone G025H7), IFN-γ–BV785 (clone 4S.B3), PD-1–PE-Cy7 (clone EH12.2H7), TIM-3–BV650 (clone F38-2E2), and TNF–BV650 (clone Mab11) from BioLegend; TCF1–AF488 (clone C63D9) from Cell Signaling; TOX–A647 (clone REA473) from Miltenyi Biotec; and CD4–PE-Cy5.5 (clone S3.5) and IL-17A–PE (clone eBio64DEC17) from Thermo Fisher Scientific.

### Peptides

Peptides corresponding to known optimal epitopes derived from CMV (pp65) and EBV (BZLF1 and EBNA-1) were purchased from Peptides & Elephants GmbH. Overlapping peptides spanning the immunogenic domains of the SARS-CoV-2 membrane (Prot_M), nucleocapsid (Prot_N), and spike proteins (Prot_S) were purchased from Miltenyi Biotec. Optimal peptides for the manufacture of HLA class I tetramers were synthesized at >95% purity by Peptides & Elephants GmbH. Lyophilized peptides were reconstituted at 10 mg/ml in DMSO and further diluted to 100 μg/ml in PBS.

### Peptide prediction

The peptide selection was made from a dataset containing all SARS-CoV-2 full length sequences from the NCBI (March 17^th^). In total, 82 different strains from 13 countries (mostly from US and China, but also including Sweden) were included. For each SARS-CoV2 amino acid sequence, the HLA peptide binding prediction method NetMHCpan-4.1 (Reynisson et al., 2020) were applied to predict conserved putative 9 mer peptide-binders to HLA-A*0201 and -B*0702. Predicted strong binders (SB) were defined as having %Rank <0.5 and weak binders (WB) <2.00 (Table S2). From 160 putative peptide binders from Spike, Envelope, Membrane, Nucleocapsid ORF3A, ORF6, ORF7a and ORF8, we identified 13 strong binders from all protein regions, that were included for tetramer generation (Table S2).

### Tetramers

HLA class I tetramers were generated as described previously (Price et al., 2005). The following specificities were used in this study: CMV A*0201 NV9 (NLVPMVATV), EBV A*0201 GL9 (GLCTLVAML), SARS-CoV-2 A*0201 AV9 (ALSKGVHFV), SARS-CoV-2 A*0201 HI9 (HLVDFQVTI), SARS-CoV-2 A*0201 KV9 (KLLEQWNLV), SARS-CoV-2 A*0201 LL9 (LLLDRLNQL), SARS-CoV-2 A*0201 LLY (LLYDANYFL), SARS-CoV-2 A*0201 SV9 (SLVKPSFYV), SARS-CoV-2 A*0201 TL9 (TLDSKTQSL), SARS-CoV-2 A*0201 VL9 (VLNDILSRL), SARS-CoV-2 A*0201 YL9 (YLQPRTFLL), SARS-CoV-2 B*0702 FI9 (FPRGQGVPI), SARS-CoV-2 B*0702 KT9 (KPRQKRTAT), SARS-CoV-2 B*0702 SA9 (SPRRARSVA), and SARS-CoV-2 B*0702 SL9 (SPRWYFYYL).

### Functional assay

PBMCs were resuspended in complete medium (RPMI 1640 supplemented with 10% FBS, 1% L-glutamine, and 1% penicillin/streptomycin) at 1 × 10^7^ cells/ml and cultured at 1 × 10^6^ cells/well in 96-well V-bottom plates (Corning) with the relevant peptides (each at 0.5 μg/ml) for 30 min prior to the addition of unconjugated anti-CD28 (clone L293) and anti-CD49d (clone L25) (each at 3 μl/ml; BD Biosciences), brefeldin A (1 μl/ml; Sigma-Aldrich), monensin (0.7 μl/ml; BD Biosciences), and anti-CD107a–PE-CF594 (clone H4A3; BD Biosciences). Negative control wells lacked peptides, and positive control wells included staphylococcal enterotoxin B (SEB; 0.5 μg/ml; Sigma-Aldrich). Cells were analyzed by flow cytometry after incubation for 8 hr at 37°C.

### Proliferation assay

PBMCs were labeled with CTV (0.5 μM; Thermo Fisher Scientific), resuspended in complete medium at 1 × 10^7^ cells/ml, and cultured at 1 × 10^6^ cells/well in 96-well U-bottom plates (Corning) with the relevant peptides (each at 0.5 μg/ml) in the presence of unconjugated anti-CD28 (clone L293) and anti-CD49d (clone L25) (each at 3 μl/ml; BD Biosciences) and IL-2 (10 IU/ml; PeproTech). Functional assays were performed as described above after incubation for 5 days at 37°C.

### AIM assay

PBMCs were resuspended in complete medium at 1 × 10^7^ cells/ml and cultured at 1 × 10^6^ cells/well in 96-well U-bottom plates (Corning) with the relevant peptides (each at 1 μg/ml) in the presence of anti-CD28 (clone L293) and anti-CD49d (clone L25) (each at 3 μl/ml; BD Biosciences). Cells were analyzed by flow cytometry after incubation for 24 hr at 37°C. The following directly conjugated monoclonal antibdodies were used to detect activation markers: anti-CD69–BUV737 (clone FN50; BD Biosciences) and anti-4-1BB–BV421 (clone 4B41; BioLegend).

### Trucount

Absolute counts from the different samples were obtained using BD Multitest™ 6-color TBNK reagents with bead-containing BD Trucount™ tubes (337166) according to manufacturer’s instructions. Samples were fixed with 2% PFA for 2 hours prior to acquiring. Absolute CD3+ cell counts were calculated using the following formula:

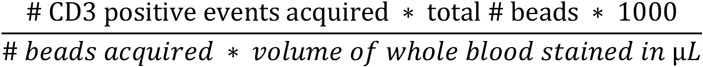

CD4^+^ and CD8^+^ counts were computed using their frequencies relative to CD3^+^ cells.

### Principal Component Analysis (PCA)

PCA were performed in Python, using scikit-learn 0.22.1. Phenotypic data obtained from flow cytometry for each cell subset was normalized using sklearn.preprocessing.StandardScaler and PCA were computed on the resulting z-scores.

### UMAP

FCS 3.0 data files were imported into FlowJo software version 10.6.0 (FlowJo LLC). All samples were compensated electronically. Dimensionality reduction was performed using the FlowJo plugin UMAP version 2.2 (FlowJo LLC). The downsample version 3.0.0 plugin and concatenation tool was used to visualize multiparametric data from up to 120,000 CD8^+^ T cells (n = 3 donors per group). The following parameters were used in these analyses: metric = euclidean, nearest neighbors = 30, and minimum distance = 0.5. Clusters of phenotypically related cells were detected using PhenoGraph version 0.2.1. The following markers were included in the cluster analysis: CCR7, CD27, CD28, CD38, CD39, CD45RA, CD95, CD127, CTLA4, CXCR5, granzyme B, Ki-67, LAG-3, PD-1, perforin, TCF-1, TIGIT, TIM-3, TOX, and 2B4. Plots were generated using Prism version 8.2.0 (GraphPad Software Inc.).

### ELISpot assay

PBMCs were rested overnight in complete medium and seeded at 2 × 10^5^ cells/well in MultiScreen HTS Filter Plates (Merck Millipore) pre-coated with anti-IFN-γ (clone 1-D1K; 15 μg/ml; Mabtech). Test wells were supplemented with overlapping peptides spanning Prot_E, Prot_N, and Prot_S (each at 2 μg/ml; Miltenyi Biotec). Negative control wells lacked peptides, and positive control wells included SEB (0.5 μg/ml; Sigma-Aldrich). Assays were incubated for 24 hr at 37°C. Plates were then washed six times with PBS (Sigma-Aldrich) and incubated for 2 hr at room temperature with biotinylated anti-IFN-γ (clone mAb-7B6-1; 1 μg/ml Mabtech). After six further washes, a 1:1,000 dilution of alkaline phosphatase-conjugated streptavidin (Mabtech) was added for 1 hr at room temperature, Plates were then washed a further six times and developed for 20 min with BCIP/NBT Substrate (Mabtech). All assays were performed in duplicate. Mean values from duplicate wells were used for data representation. Spots were counted using an automated ELISpot Reader System (Autoimmun Diagnostika GmbH).

### Serology

Serum samples from all donors were barcoded and dispatched to Clinical Microbiology, Karolinska University Laboratory for serology assessment. SARS-CoV-2-specific antibodies were detected using both the iFLASH Anti-SARS-CoV-2 IgG chemiluminescent microparticle immunoassay against the nucleocapsid and envelope proteins (Shenzhen Yhlo Biotech Co. Ltd.) as well as the LIAISON SARS-CoV-2 IgG fully automated indirect chemiluminescent immunoassay serology assay against the S1 and S2 (spike) proteins (DiaSorin). The assays produced highly concordant results (Figure S8A) and have both been shown to generate satisfactory diagnostic performance as serological SARS-CoV-2 assays (Plebani et al., 2020). An individual was considered seropositive if one of the two methods generated a positive result. All assays were performed by trained employees at the clinical laboratory according to the respective manufacturer standard procedures.

### Statistics

Statistical analyses were performed using R studio or Prism version 7.0 (GraphPad Software Inc.). Phenotypic relationships within multivariate data sets were visualized using FlowJo software version 10.6.1 (FlowJo LLC). Differences between unmatched groups were compared using an unpaired t-test or the Mann-Whitney *U* test, and differences between matched groups were compared using a paired t-test or the Wilcoxon signed-rank test. Correlations were assessed using the Pearson correlation or the Spearman rank correlation. Non-parametric tests were used if the data were not distributed normally according to the Shapiro-Wilk normality test.

## ACKNOWLEDGEMENTS

We express our gratitude to all donors, health care personnel, study coordinators, administrators, and laboratory managers involved in this work.

## AUTHOR CONTRIBUTIONS

H.G.L., S.A., and M.B. conceived the project; T.S., A.P.P., O.R.B., J.B.G., S.L.L., S.M., D.J.W., T.K., J.E., T.P., D.A.P., and M.B. designed and performed experiments; T.S., A.P.P., O.R.B., J.B.G., and M.B. analyzed data; K.S., A.O., G.B., E.F., O.R, L.I.E, A.S., T.A., J.A., M.N., J.K., S.G.R., N.K.B., J.K.S., D.A.P., H.G.L., S.A., and M.B. provided critical resources; D.A.P. and M.B. supervised experiments; T.S., A.P.P., O.R.B., and M.B. drafted the manuscript; D.A.P. and M.B. edited the manuscript. All authors, including the Karolinska COVID-19 Study Group, contributed intellectually and approved the manuscript.

## DATA AND MATERIALS AVAILABILITY

Data are available on request from the corresponding author.

## Karolinska COVID-19 Study Group

^1^Mira Akber, ^2^Soo Aleman, ^1^Lena Berglin, ^1^Helena Bergsten, ^1^Niklas K Björkström, ^1^Susanna Brighenti, ^1^Demi Brownlie, ^1^Marcus Buggert, ^1^Marta Butrym, ^1^Benedict Chambers, ^1^Puran Chen, ^1^Martin Cornillet, ^4^Jonathan Grip, ^1^Angelica Cuapio Gomez, ^2^Lena Dillner, ^1^Jean-Baptiste Gorin, ^1^Isabel Diaz Lozano, ^1^Majda Dzidic, ^1^Johanna Emgård, ^3^Lars I Eriksson, ^1^Malin Flodström Tullberg, ^2^Anna Färnert, ^2^Hedvig Glans, ^1^Sara Gredmark Russ, ^1^Alvaro Haroun-Izquierdo, ^1^Elizabeth Henriksson, ^1^Laura Hertwig, ^2^Habiba Kamal, ^1^Tobias Kammann, ^1^Jonas Klingstrom, ^1^Efthymia Kokkinou, ^1^Egle Kvedaraite, ^1^Hans-Gustaf Ljunggren, ^1^Marco Loreti, ^1^Magalini Lourda, ^1^Kimia Maleki, ^1^Karl-Johan Malmberg, ^1^Christopher Maucourant, ^1^Jakob Michaelsson, ^1^Jenny Mjösberg, ^1^Kirsten Moll, ^1^ Jagadeeswara R. Muvva, ^3^Johan Mårtensson, ^2^Pontus Nauclér, ^1^Anna Norrby-Teglund, ^2^Annika Olsson, ^1^Laura Palma Medina, ^1^Tiphaine Parrot, ^3^Björn Persson, ^1^André Perez-Potti, ^1^Lena Radler, ^1^Emma Ringqvist, ^1^Olga Rivera-Ballesteros, ^4^Olav Rooyackers, ^1^Johan K. Sandberg, ^1^John Tyler Sandberg, ^1^Takuya Sekine, ^1^Ebba Sohlberg, ^1^Tea Soini, ^2^Kristoffer Strålin, ^2^Anders Sönnerborg, ^1^Mattias Svensson, ^1^Janne Tynell, ^1^Renata Varnaite, ^1^Andreas Von Kries, ^5^Christian Unge, ^1^David J. Wullimann.

^1^Center for Infectious Medicine, Department of Medicine Huddinge, Karolinska Institutet, Karolinska University Hospital, Stockholm, Sweden

^2^Division of Infectious Diseases and Dermatology, Karolinska University Hospital and Department of Medicine Huddinge, Stockholm, Sweden

^3^Department of Physiology and Pharmacology, Karolinska Institutet and Function Perioperative Medicine and Intensive Care, Karolinska University Hospital, Stockholm, Sweden

^4^Department of Physiology and Pharmacology, Karolinska Institutet and Function Perioperative Medicine and Intensive Care, Karolinska University Hospital, Stockholm, Sweden

^5^Department of Emergency Medicine, Karolinska University Hospital, Stockholm, Sweden

---

**Table S1.** Donor characteristics.

|  | Acute<br>severe<br>(n = 17) | Acute<br>moderate<br>(n = 10) | Severe<br>convalescent<br>(n = 26) | Mild convalescent<br>(n = 40) | Exposed relatives<br>(n = 30) | 2020<br>blood donors<br>(n = 55) | 2019<br>blood donors<br>(n = 25) |
| --- | --- | --- | --- | --- | --- | --- | --- |
| Age<br>(median, years) | 58 | 52 | 51 | 51 | 42 | 49 | n/d |
| Gender<br>(% male/female) | 82/18 | 70/30 | 88/12 | 45/55 | 33/67 | n/d | n/d |
| Travel | n/d | n/d | 33% | 85%<br>(79% Italy) | 33%<br>(100% Italy) | n/d | n/d |
| BMI<br>(median) | 32 | 28 | 28 | 24 | n/d | n/d | n/d |
| Smoker | 44%<br>(past/now) | Unclear | 38%<br>(past/now) | 23%<br>(past/now) | n/d | n/d | n/d |
| Diabetes | 29% | 30% | 35% | 10% | 0% | n/d | n/d |
| Hypertension | 35% | 20% | 42% | 15% | 0.7% | n/d | n/d |
| Other chronic<br>diseases | 71% | 50% | 19% | 15% | 17% | n/d | n/d |
| PCR<br>positivity | 100% | 100% | 100% | 100% | n/d | n/d | n/d |
| Viremia at time<br>of sampling | 41% | 40% | n/d | n/d | n/d | n/d | n/d |
| Asymptomatic | 0% | 0% | 0% | 5% | 21% | n/d | n/d |
| Outcome | 76%<br>alive | 100%<br>alive | 100%<br>alive | 100%<br>alive | 100%<br>alive | n/d | n/d |
| Antibody<br>positivity | 82% | 50% | 100% | 85% | 64% | 7% | n/d |

BMI: body mass index (18.5-24.5 normal; 25-29.9 overweight; >30 obesity); n/d: not determined.

**Table S2.**
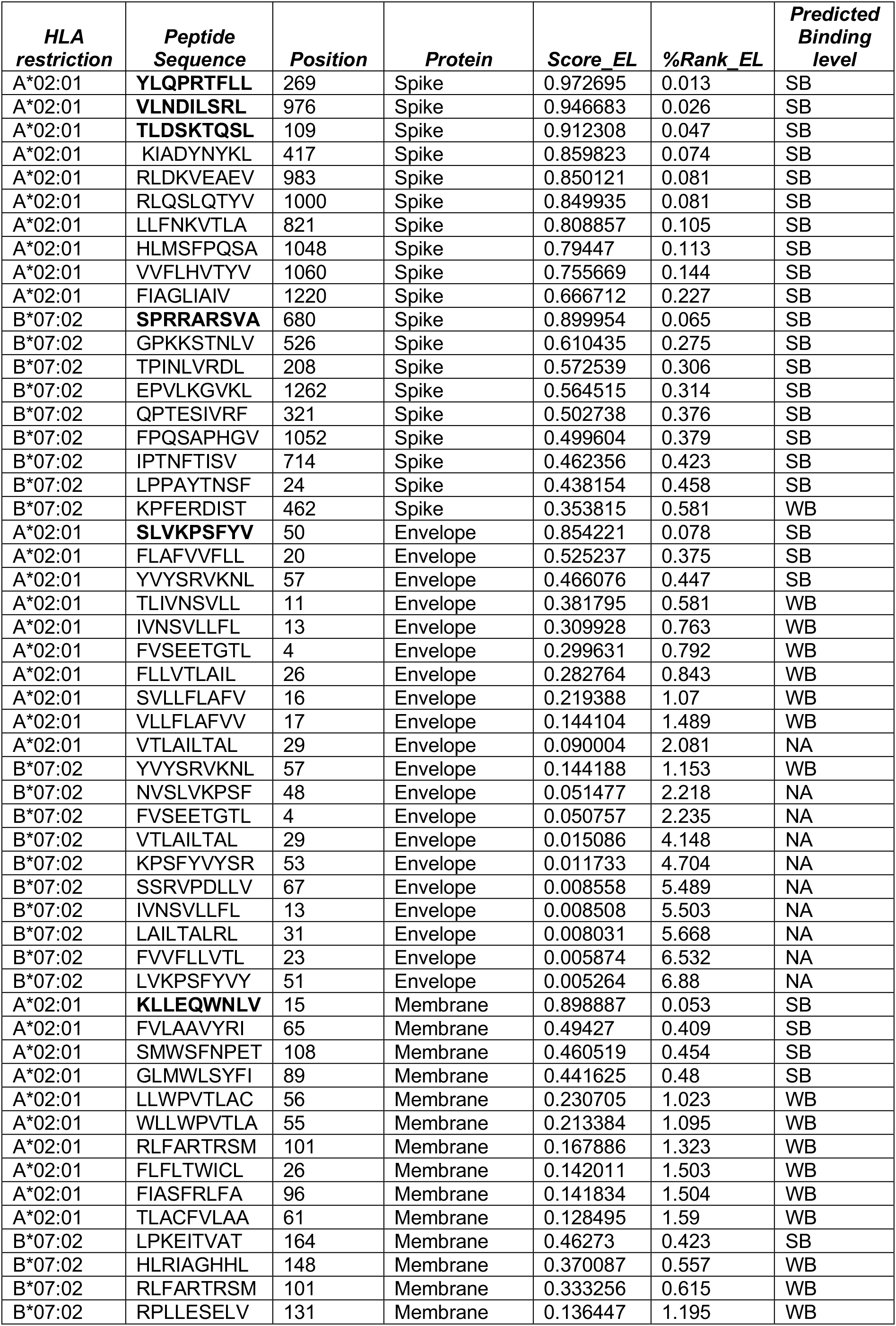
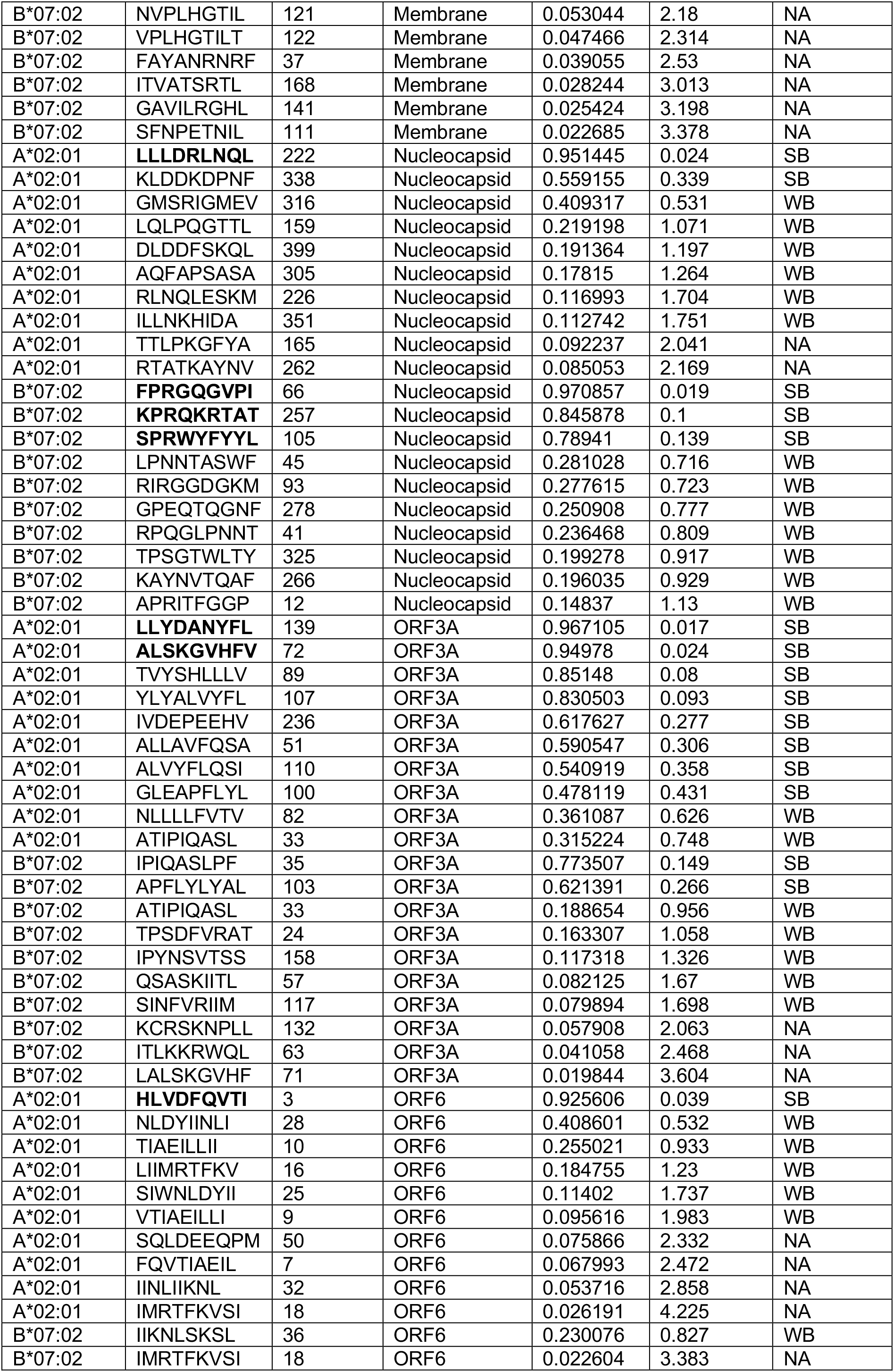
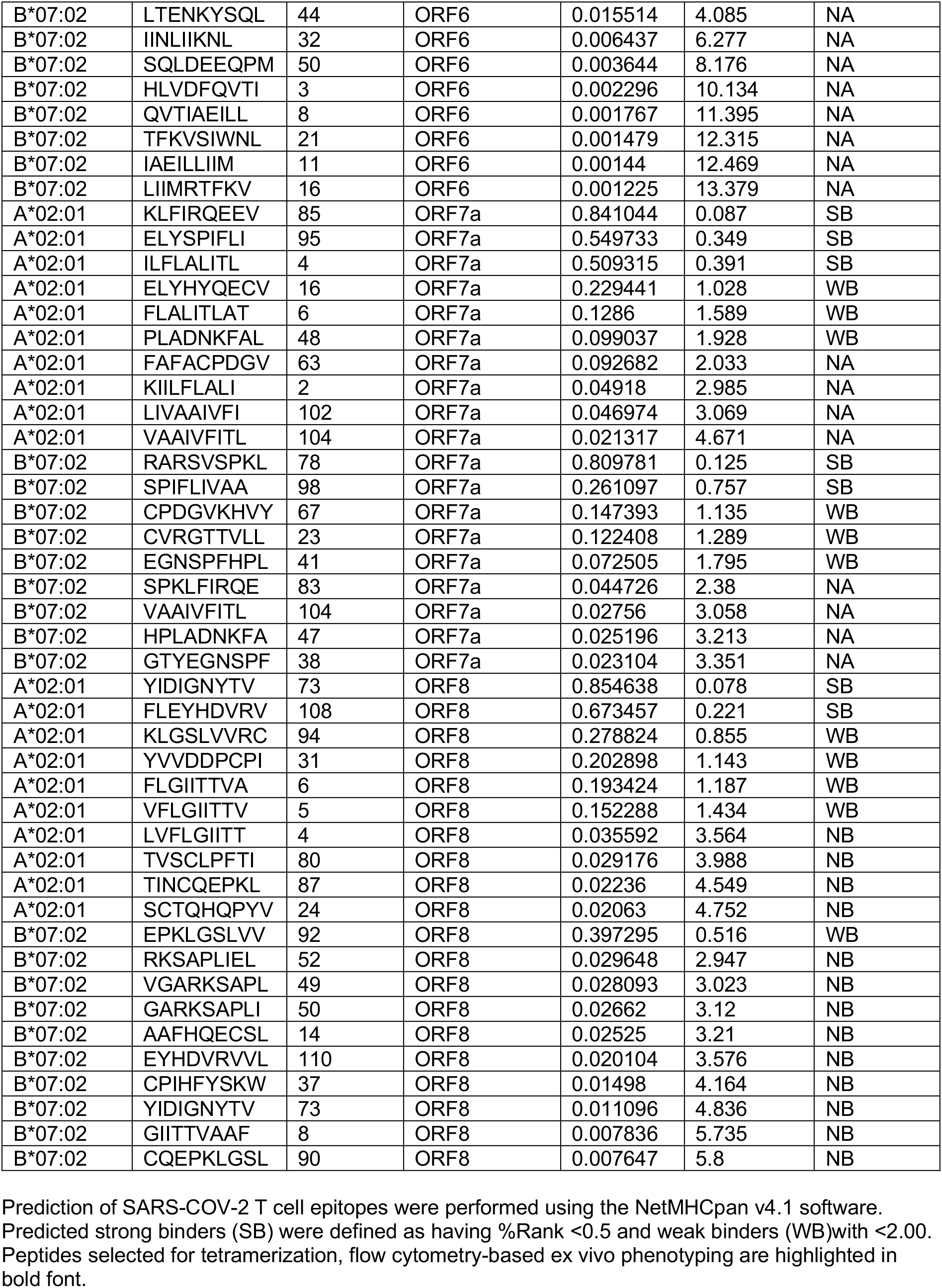
Predicted SARS-COV-2 T cell epitopes and their HLA binding affinity

