## Supplemental info for "Robust T cell immunity in convalescent individuals with asymptomatic or mild COVID-19"

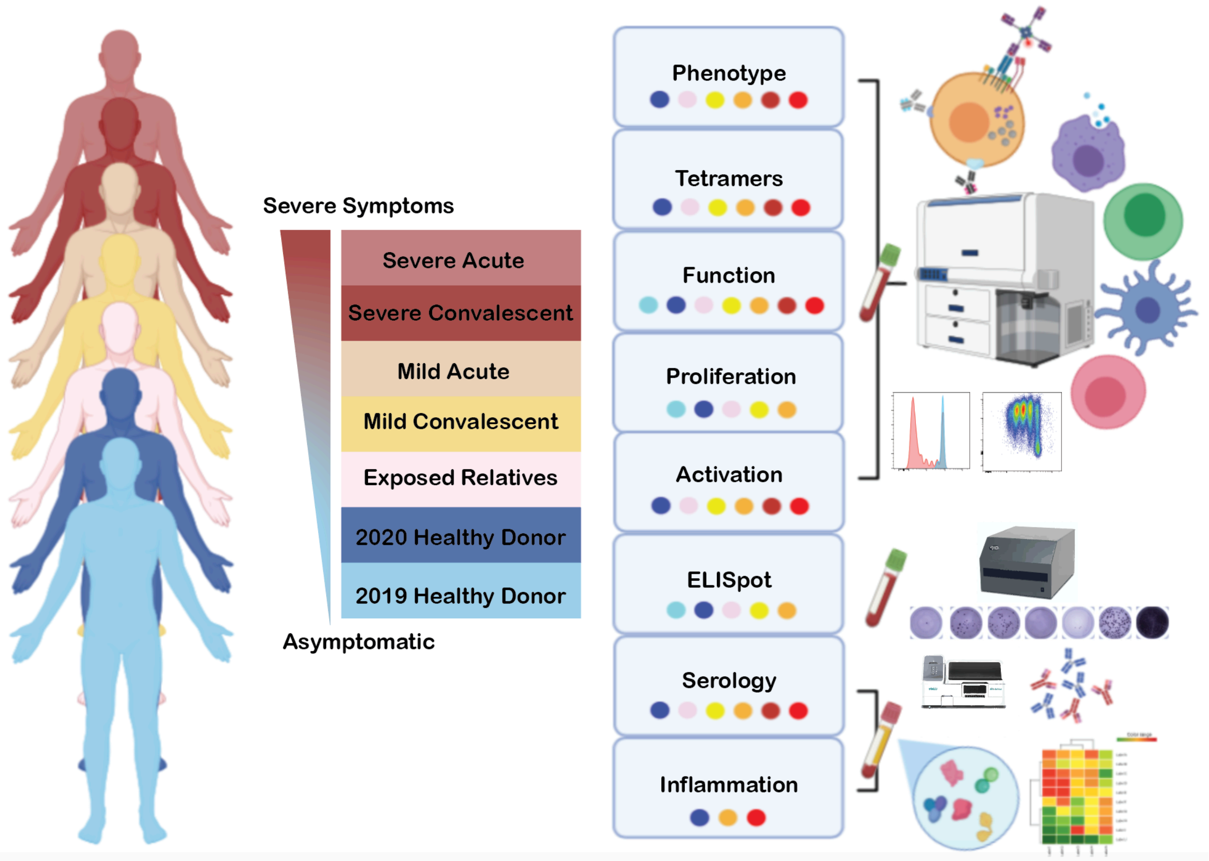


**Figure S1. Cohort definition and workflow.** Overview of donor groups and experimental assays. Colored dots indicate the groups included in each set of assays. Related to Figures 1, 2, 3, and 4.

Figure S2

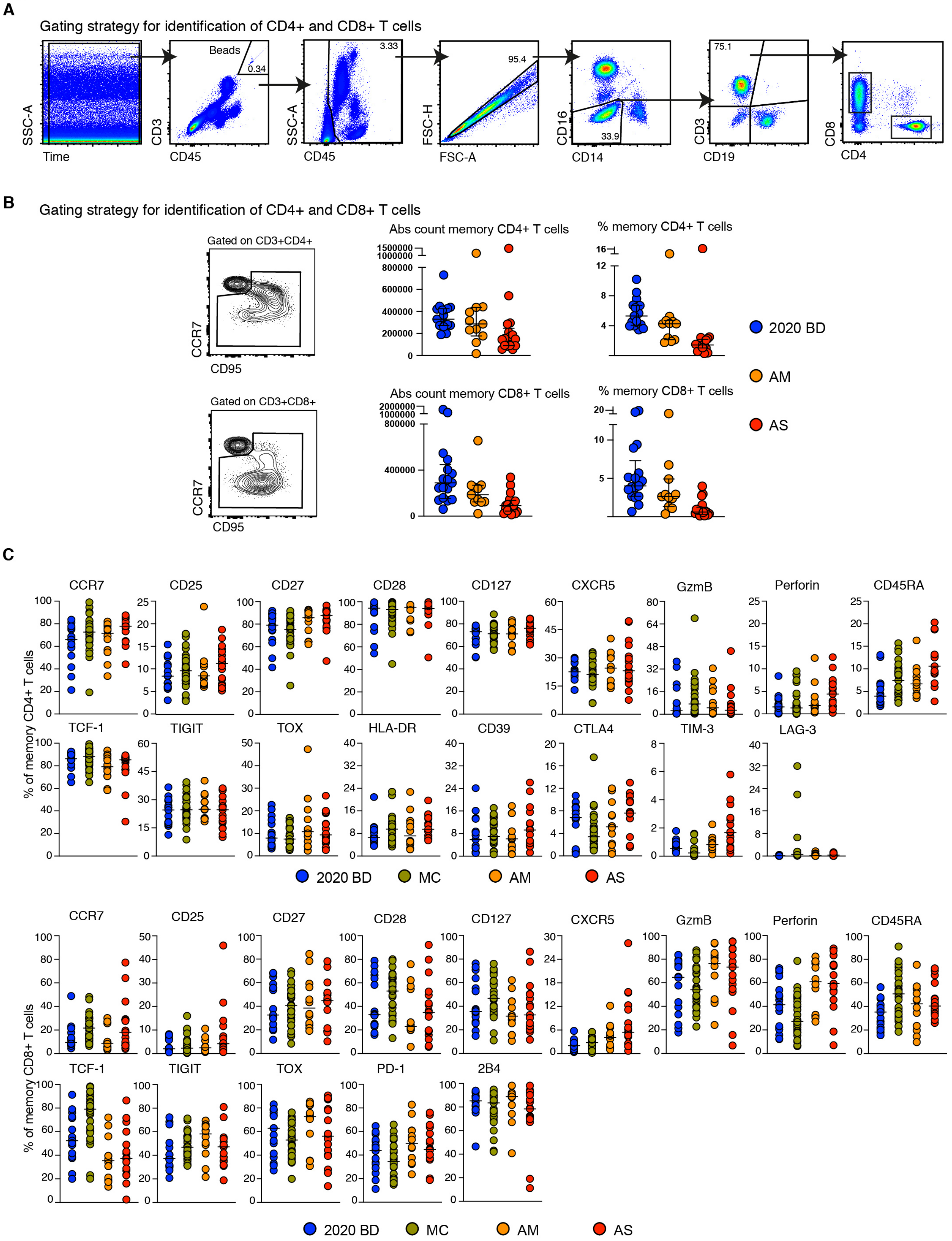


**Figure S2. Quantification and characterization of CD4$^{+}$ and CD8$^{+}$ T cells in COVID-19.** (**A**) Flow cytometric gating strategy for the identification and quantification of CD4$^{+}$ and CD8$^{+}$ T cells. (**B**) Left: flow cytometric gating strategy for the identification and quantification of memory $CD4^+$ and $CD8^+$ T cells. Right: dot plots summarizing the absolute numbers and relative frequencies of memory $CD4^+$ and $CD8^+$ T cells by group. Each dot represents one donor. Data are shown as median ± IQR. 2020 BD: healthy blood donors from 2020. AM: patients with acute moderate COVID-19. AS: patients with acute severe COVID-19. (**C**) Dot plots showing the expression frequencies of phenotypic markers among memory $CD4^+$ and $CD8^+$ T cells by group. Each dot represents one donor. Bars indicate median values. 2020 BD: healthy blood donors from 2020. MC: individuals in the convalescent phase after asymptomatic/mild COVID-19. AM: patients with acute moderate COVID-19. AS: patients with acute severe COVID-19. Related to Figure 1.

Figure S3

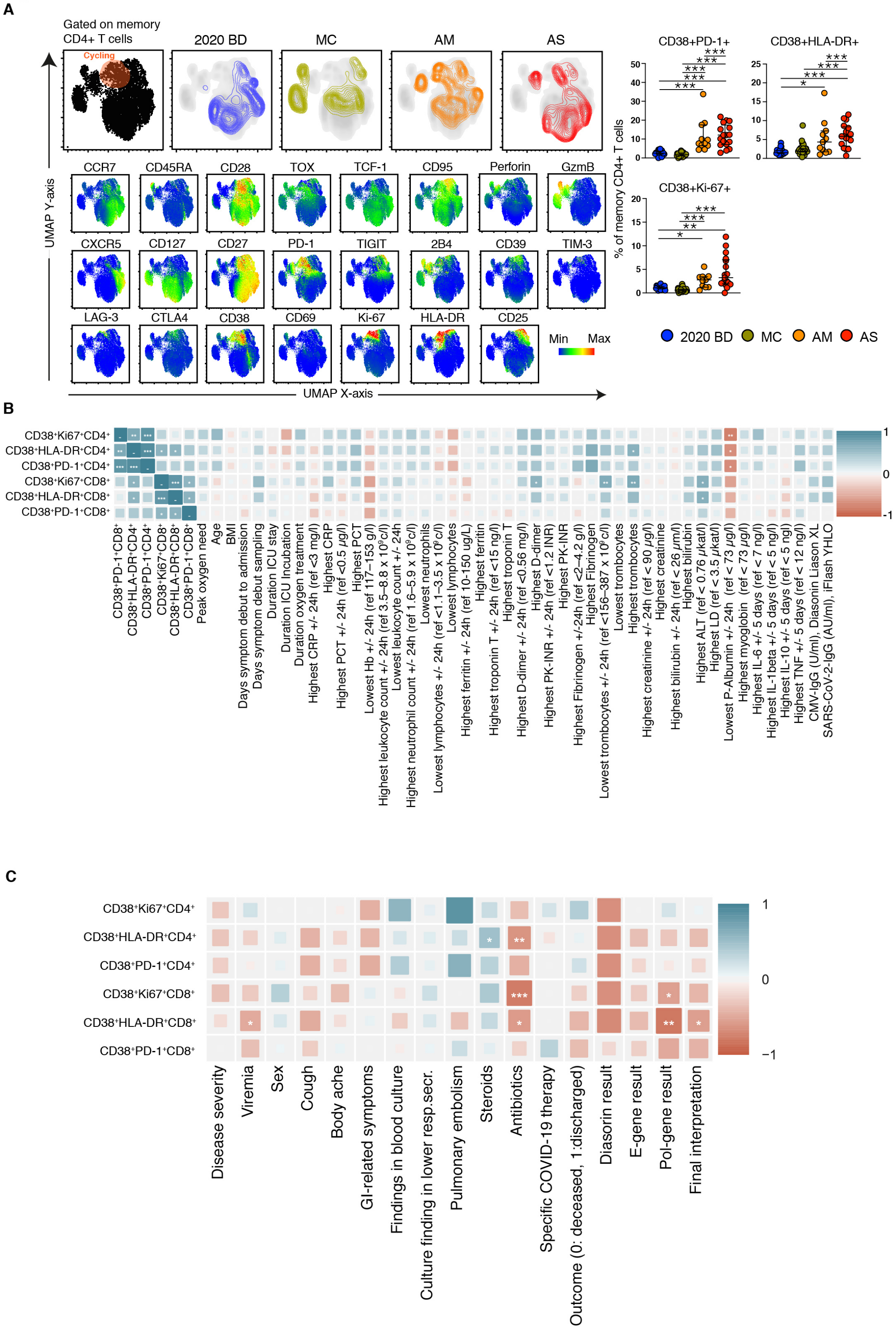
A
Gated on memory CD4+ T cells
Cycling
2020 BD
MC
AM
AS
CCR7
CD45RA
CD28
TOX
TCF-1
CD95
Perforin
GzmB
CXCR5
CD127
CD27
PD-1
TIGIT
2B4
CD39
TIM-3
LAG-3
CTLA4
CD38
CD69
Ki-67
HLA-DR
CD25
Min
Max
UMAP Y-axis
UMAP X-axis
CD38+PD-1+
CD38+HLA-DR+
CD38+Ki-67+
% of memory CD4+ T cells
2020 BD
MC
AM
AS
B
CD38⁺Ki67⁺CD4⁺
CD38⁺HLA-DR⁺CD4⁺
CD38⁺PD-1⁺CD4⁺
CD38⁺Ki67⁺CD8⁺
CD38⁺HLA-DR⁺CD8⁺
CD38⁺PD-1⁺CD8⁺
Peak oxygen need
Age
BMI
Days symptom debut to admission
Days symptom debut sampling
Duration ICU stay
Duration ICU Incubation
Duration oxygen treatment
Highest CRP +/- 24h (ref <3 mg/l)
Highest CRP
Highest PCT +/- 24h (ref <0.5 µg/l)
Highest PCT
Lowest Hb +/- 24h (ref 117–153 g/l)
Highest leukocyte count +/- 24h (ref 3.5–8.8 x 10⁹c/l)
Lowest leukocyte count +/- 24h
Highest neutrophil count +/- 24h (ref 1.6–5.9 x 10⁹c/l)
Lowest neutrophils
Lowest lymphocytes +/- 24h (ref <1.1–3.5 x 10⁹c/l)
Lowest lymphocytes
Highest ferritin +/- 24h (ref 10-150 ug/L)
Highest ferritin
Highest troponin T +/- 24h (ref <15 ng/l)
Highest troponin T
Highest D-dimer +/- 24h (ref <0.56 mg/l)
Highest D-dimer
Highest PK-INR +/- 24h (ref <1.2 INR)
Highest PK-INR
Highest Fibrinogen +/-24h (ref <2–4.2 g/l)
Highest Fibrinogen
Lowest trombocytes +/- 24h (ref <156–387 x 10⁹c/l)
Lowest trombocytes
Highest trombocytes
Highest creatinine +/- 24h (ref < 90 µg/l)
Highest creatinine
Highest bilirubin +/- 24h (ref < 26 µm/l)
Highest bilirubin
Highest ALT (ref < 0.76 µkat/l)
Highest LD (ref < 3.5 µkat/l)
Lowest P-Albumin +/- 24h (ref < 73 µg/l)
Highest myoglobin (ref < 73 µg/l)
Highest IL-6 +/- 5 days (ref < 7 ng/l)
Highest IL-1beta +/- 5 days (ref < 5 ng/l)
Highest IL-10 +/- 5 days (ref < 5 ng/l)
Highest TNF +/- 5 days (ref < 12 ng/l)
CMV-IgG (U/ml), Diasonin Liason XL
SARS-CoV-2-IgG (AU/ml), iFlash YHLO
C
CD38⁺Ki67⁺CD4⁺
CD38⁺HLA-DR⁺CD4⁺
CD38⁺PD-1⁺CD4⁺
CD38⁺Ki67⁺CD8⁺
CD38⁺HLA-DR⁺CD8⁺
CD38⁺PD-1⁺CD8⁺
Disease severity
Viremia
Sex
Cough
Body ache
GI-related symptoms
Findings in blood culture
Culture finding in lower resp.secr.
Pulmonary embolism
Steroids
Antibiotics
Specific COVID-19 therapy
Outcome (0: deceased, 1:discharged)
Diasorin result
E-gene result
Pol-gene result
Final interpretation

**Figure S3. UMAP clustering of memory $CD4^+$ T cells and correlative analyses of immune activation phenotypes versus clinical parameters in acute COVID-19.** (**A**) Top left: UMAP plots showing the clustering of memory $CD4^+$ T cell by group in relation to all memory $CD4^+$ T cells (left). Bottom left: UMAP plots showing the expression of individual markers (n = 3 donors per group). Right: dot plots summarizing the expression frequencies of activation/cycling markers among memory $CD4^+$ T cells by group. Each dot represents one donor. Data are shown as median ± IQR. 2020 BD: healthy blood donors from 2020. MC: individuals in the convalescent phase after asymptomatic/mild COVID-19. AM: patients with acute moderate COVID-19. AS: patients with acute severe COVID-19. (**B** and **C**) Heatmaps summarizing the pairwise correlations between phenotypically defined subpopulations of memory $CD4^+$ or $CD8^+$ T cells and various clinical parameters in patients with acute COVID-19. $^*P < 0.05$, $^{**}P < 0.01$, $^{***}P < 0.001$. Related to Figure 1.

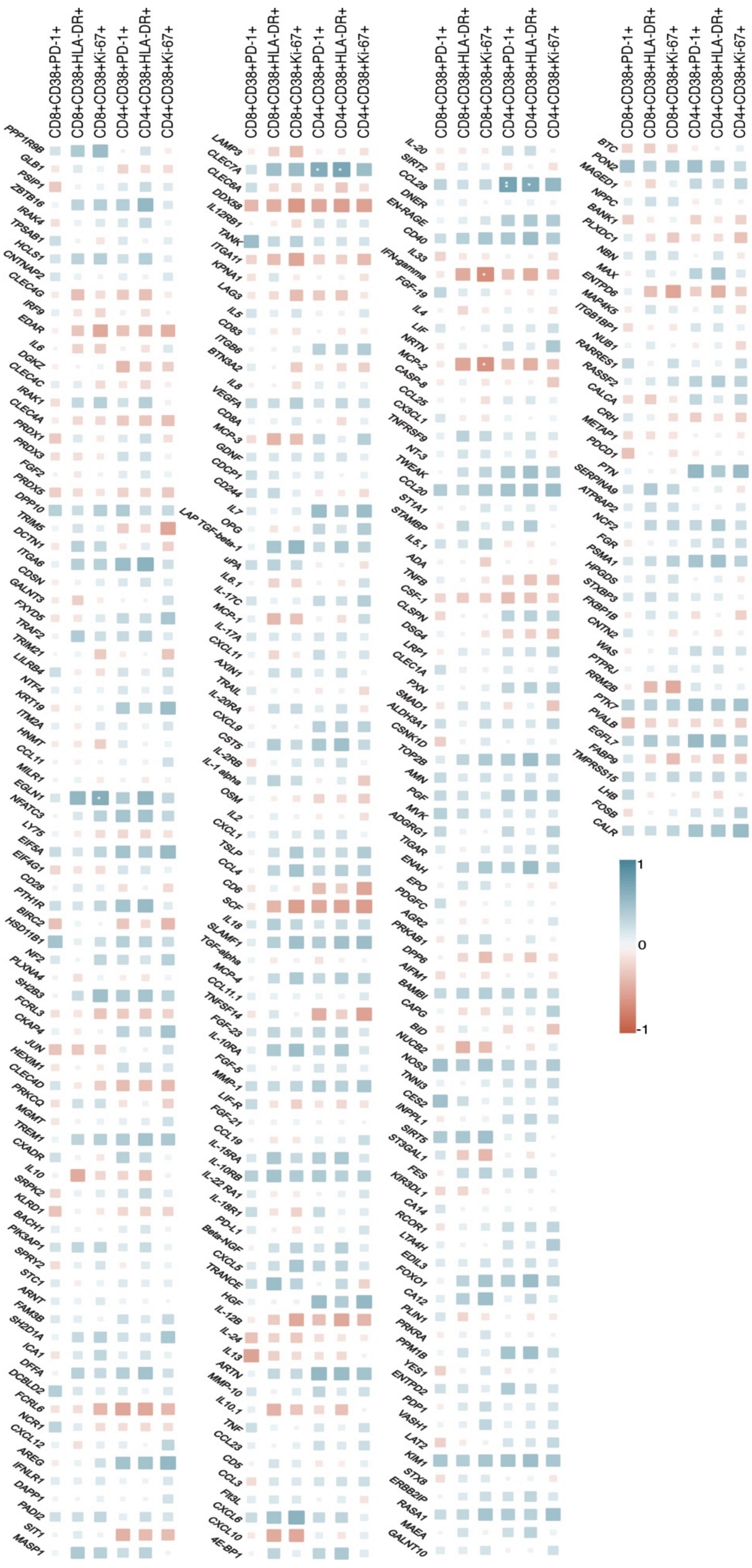
CD8+CD38+PD-1+
CD8+CD38+HLA-DR+
CD8+CD38+Ki-67+
CD4+CD38+PD-1+
CD4+CD38+HLA-DR+
CD4+CD38+Ki-67+
PPP1R9B
GLB1
PSIP1
ZBTB16
IRAK4
TPSAB1
HCLS1
CNTNAP2
CLEC4G
IRF9
EDAR
IL6
DGKZ
CLEC4C
IRAK1
CLEC4A
PRDX1
PRDX3
FGF2
PRDX5
DPP10
TRIM5
DCTN1
ITGA6
CDSN
GALNT3
FXYD5
TRAF2
TRIM21
LILRB4
NTF4
KRT19
ITM2A
HNMT
CCL11
MILR1
EGLN1
NFATC3
LY75
EIF5A
EIF4G1
CD28
PTH1R
BIRC2
HSD11B1
NF2
PLXNA4
SH2B3
FCRL3
CKAP4
JUN
HEXIM1
CLEC4D
PRKCQ
MGMT
TREM1
CXADR
IL10
SRPK2
KLRD1
BACH1
PIK3AP1
SPRY2
STC1
ARNT
FAM3B
SH2D1A
ICA1
DFFA
DCBLD2
FCRL6
NCR1
CXCL12
AREG
IFNLR1
DAPP1
PADI2
SIT1
MASP1
CD8+CD38+PD-1+
CD8+CD38+HLA-DR+
CD8+CD38+Ki-67+
CD4+CD38+PD-1+
CD4+CD38+HLA-DR+
CD4+CD38+Ki-67+
LAMP3
CLEC7A
CLEC6A
DDX58
IL12RB1
TANK
ITGA11
KPNA1
LAG3
IL5
CD83
ITGB6
BTN3A2
IL8
VEGFA
CD8A
MCP-3
GDNF
CDCP1
CD244
IL7
OPG
LAP TGF-beta-1
uPA
IL6.1
IL-17C
MCP-1
IL-17A
CXCL11
AXIN1
TRAIL
IL-20RA
CXCL9
CST5
IL-2RB
IL-1 alpha
OSM
IL2
CXCL1
TSLP
CCL4
CD6
SCF
IL18
SLAMF1
TGF-alpha
MCP-4
CCL11.1
TNFSF14
FGF-23
IL-10RA
FGF-5
MMP-1
LIF-R
FGF-21
CCL19
IL-15RA
IL-10RB
IL-22 RA1
IL-18R1
PD-L1
Beta-NGF
CXCL5
TRANCE
HGF
IL-12B
IL-24
IL13
ARTN
MMP-10
IL10.1
TNF
CCL23
CD5
CCL3
Flt3L
CXCL6
CXCL10
4E-BP1
CD8+CD38+PD-1+
CD8+CD38+HLA-DR+
CD8+CD38+Ki-67+
CD4+CD38+PD-1+
CD4+CD38+HLA-DR+
CD4+CD38+Ki-67+
IL-20
SIRT2
CCL28
DNER
EN-RAGE
CD40
IL33
IFN-gamma
FGF-19
IL4
LIF
NRTN
MCP-2
CASP-8
CCL25
CX3CL1
TNFRSF9
NT-3
TWEAK
CCL20
ST1A1
STAMBP
IL5.1
ADA
TNFB
CSF-1
CLSPN
DSG4
LRP1
CLEC1A
PXN
SMAD1
ALDH3A1
CSNK1D
TOP2B
AMN
PGF
MVK
ADGRG1
TIGAR
ENAH
EPO
PDGFC
AGR2
PRKAB1
DPP6
AIFM1
BAMBI
CAPG
BID
NUCB2
NOS3
TNNI3
CES2
INPPL1
SIRT5
ST3GAL1
FES
KIR3DL1
CA14
RCOR1
LTA4H
EDIL3
FOXO1
CA12
PLIN1
PRKRA
PPM1B
YES1
ENTPD2
PDP1
VASH1
LAT2
KIM1
STX8
ERBB2IP
RASA1
MAEA
GALNT10
CD8+CD38+PD-1+
CD8+CD38+HLA-DR+
CD8+CD38+Ki-67+
CD4+CD38+PD-1+
CD4+CD38+HLA-DR+
CD4+CD38+Ki-67+
BTC
PON2
MAGED1
NPPC
BANK1
PLXDC1
NBN
MAX
ENTPD6
MAP4K5
ITGB1BP1
NUB1
RARRES1
RASSF2
CALCA
CRH
METAP1
PDCD1
PTN
SERPINA9
ATP6AP2
NCF2
FGR
PSMA1
HPGDS
STXBP3
FKBP1B
CNTN2
WAS
PTPRJ
RRM2B
PTK7
PVALB
EGFL7
FABP9
TMPRSS15
LHB
FOSB
CALR
1
0
-1

**Figure S4. Correlative analyses of immune activation phenotypes versus soluble factor measurements in acute COVID-19.** Heatmap summarizing the pairwise correlations between phenotypically defined subpopulations of memory $CD4^+$ and $CD8^+$ T cells and the relative concentrations of various soluble factors in the peripheral blood of patients with acute COVID-19. **P* < 0.05, ***P* < 0.01. Related to Figure 1.

Figure S5

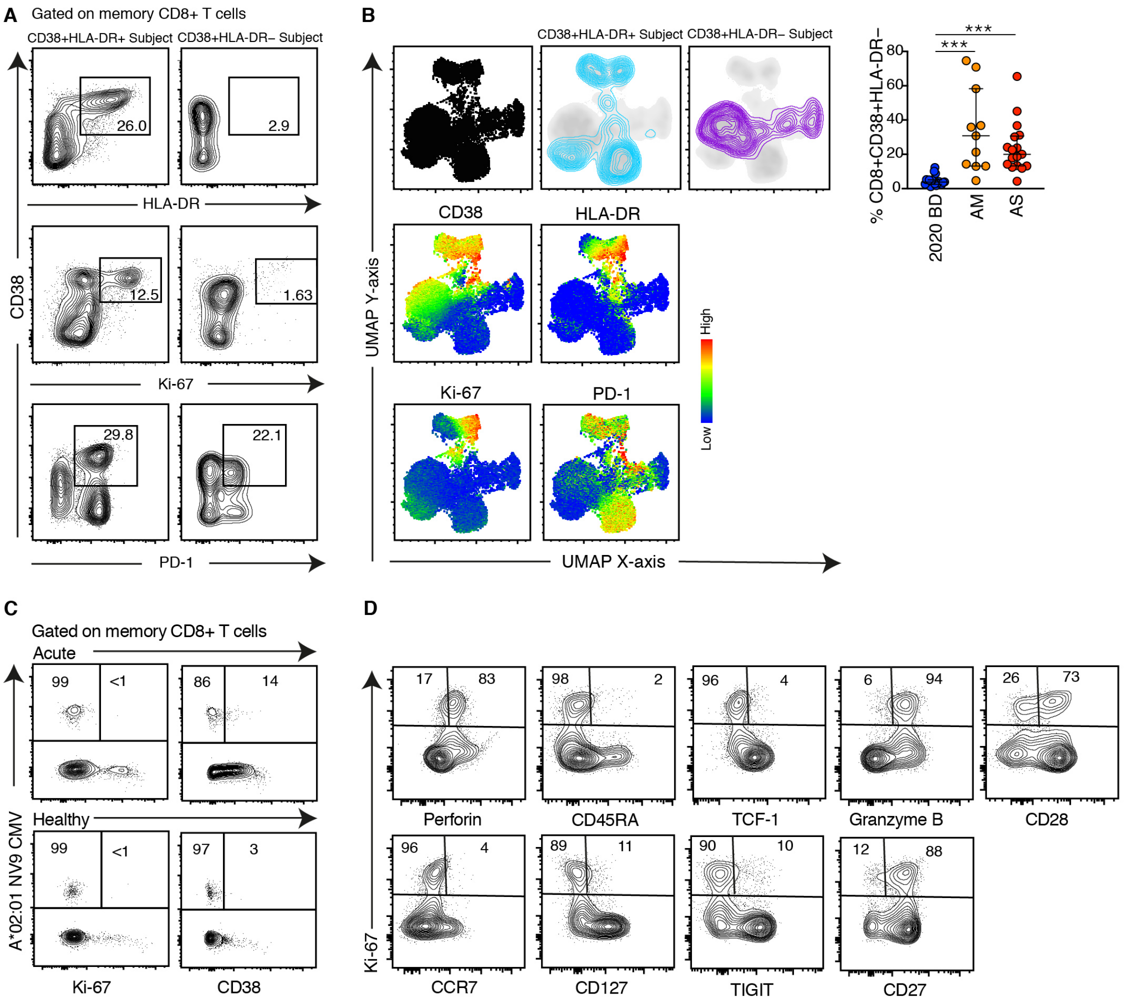


**Figure S5. Immune activation patterns in acute COVID-19.** (**A**) Representative flow cytometry plots showing the expression of activation/cycling markers among memory $CD8^+$ T cells in patients with acute severe COVID-19. Numbers indicate percentages in the drawn gates. (**B**) Top left: UMAP plots showing the clustering of memory $CD8^+$ T cells by phenotype in relation to all memory $CD8^+$ T cells (left). Bottom left: UMAP plots showing the expression of individual markers (n = 1 donor per group). Right: dot plots summarizing the frequencies of $CD38^+$ $HLA\text{-}DR^-$ memory $CD8^+$ T cells by group. Each dot represents one donor. Data are shown as median ± IQR. 2020 BD: healthy blood donors from 2020. AM: patients with acute moderate COVID-19. AS: patients with acute severe COVID-19. **(C)** Representative flow cytometry plots showing the expression of activation/cycling markers among CMV-specific memory $CD8^+$ T cells in a healthy control and a patient with acute severe COVID-19. Numbers indicate percentages in the drawn gates. **(D)** Representative flow cytometry plots illustrating phenotype of $Ki\text{-}67^+$ memory $CD8^+$ T cells in a patient with acute severe COVID-19. Numbers indicate percentages in the drawn gates. ***$P < 0.001$. Related to Figure 2.

Figure S6

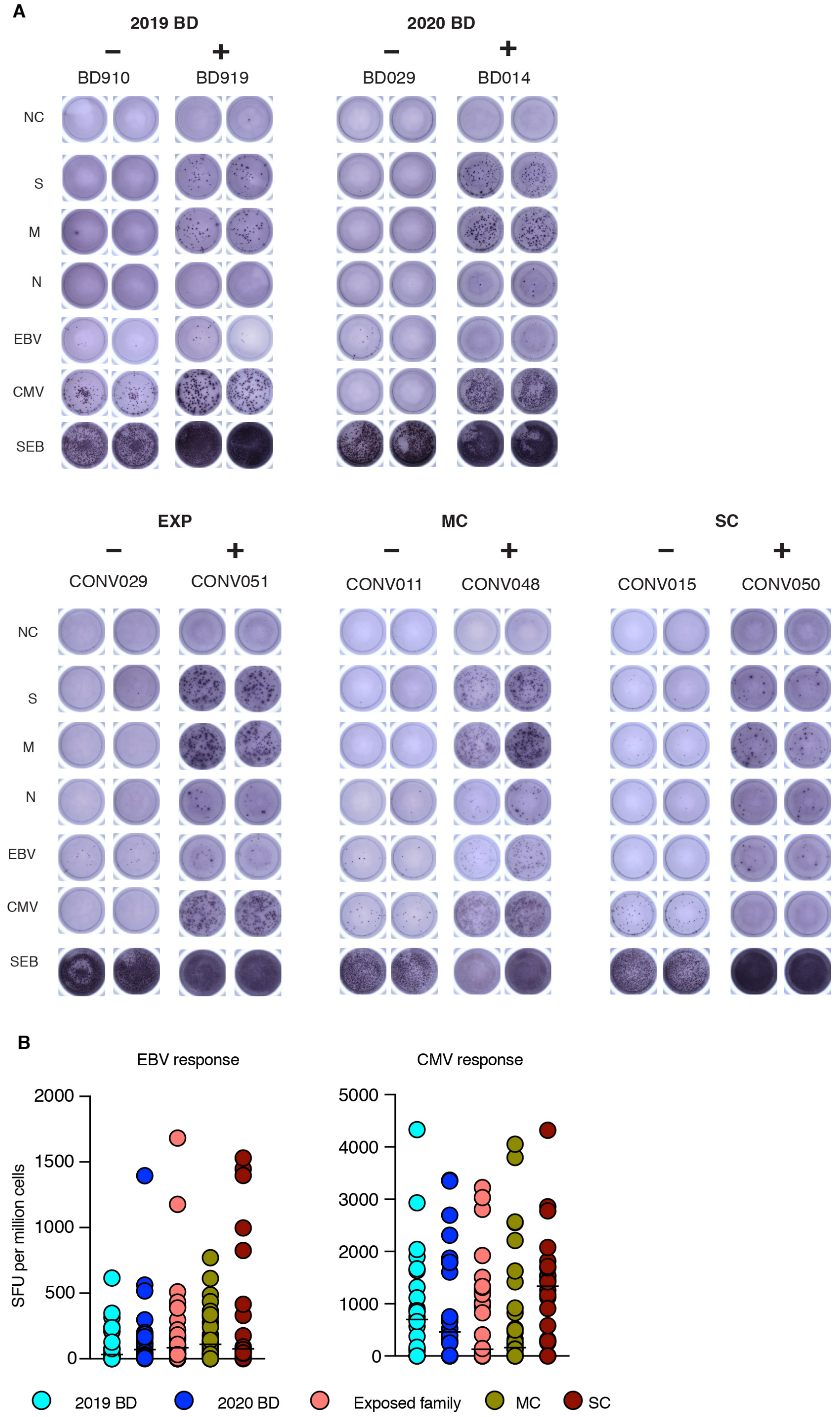


**Figure S6. Quantification of functional T cell reactivity in COVID-19.** (**A**) Representative images showing the detection of IFN-γ-producing cells responding to overlapping peptides spanning the immunogenic domains of the SARS-CoV-2 membrane (M), nucleocapsid (N), and spike proteins (S) by group (ELISpot assays). NC: negative control. EBV: Epstein-Barr virus. CMV: cytomegalovirus. SEB: staphylococcal enterotoxin B. (**B**) Dot plots summarizing the frequencies of IFN-γ-producing cells responding to optimal peptide epitopes derived from EBV BZLF1 and EBNA-1 (left) or CMV pp65 (right) by group (ELISpot assays). Each dot represents the average SFU in one donor. Bars indicate median values. No significant differences were detected among groups for any specificity. 2019 BD: healthy blood donors from 2019. 2020 BD: healthy blood donors from 2020. Exp: exposed family members. MC: individuals in the convalescent phase after asymptomatic/mild COVID-19. SC: individuals in the convalescent phase after severe COVID-19. SFU: spot-forming unit. Related to Figure 3.

Figure S7

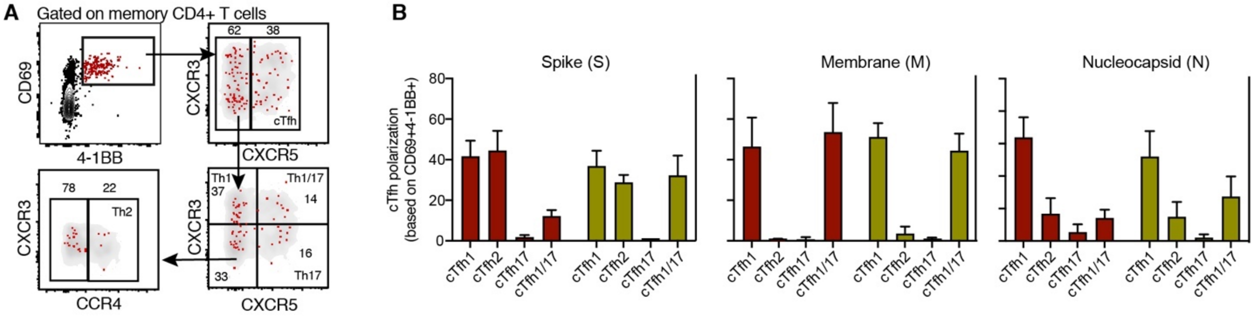


**Figure S7. Functional polarization of SARS-CoV-2-specific memory CD4$^+$ T cells in COVID-19. (A)** Representative flow cytometry plots showing the identification of memory CD4$^+$ T cells responding to overlapping peptides spanning the immunogenic domains of the SARS-CoV-2 nucleocapsid protein by subset (AIM assay). Subsets were defined as CXCR5$^+$ (cTfh), CCR4$^-$ CCR6$^-$ CXCR3$^+$ CXCR5$^-$ (Th1), CCR4$^+$ CCR6$^-$ CXCR3$^-$ CXCR5$^-$ (Th2), CCR4$^-$ CCR6$^+$ CXCR3$^-$ CXCR5$^-$ (Th17), CCR4$^-$ CCR6$^+$ CXCR3$^+$ CXCR5$^-$ (Th1/17), and CCR4$^-$ CCR6$^-$ CXCR3$^-$ CXCR5$^-$ (non-Th1/2/17). **(B)** Cumulative data of cTh polarization observed against various SARS-CoV-2 proteins. Related to Figure 3.

Figure S8

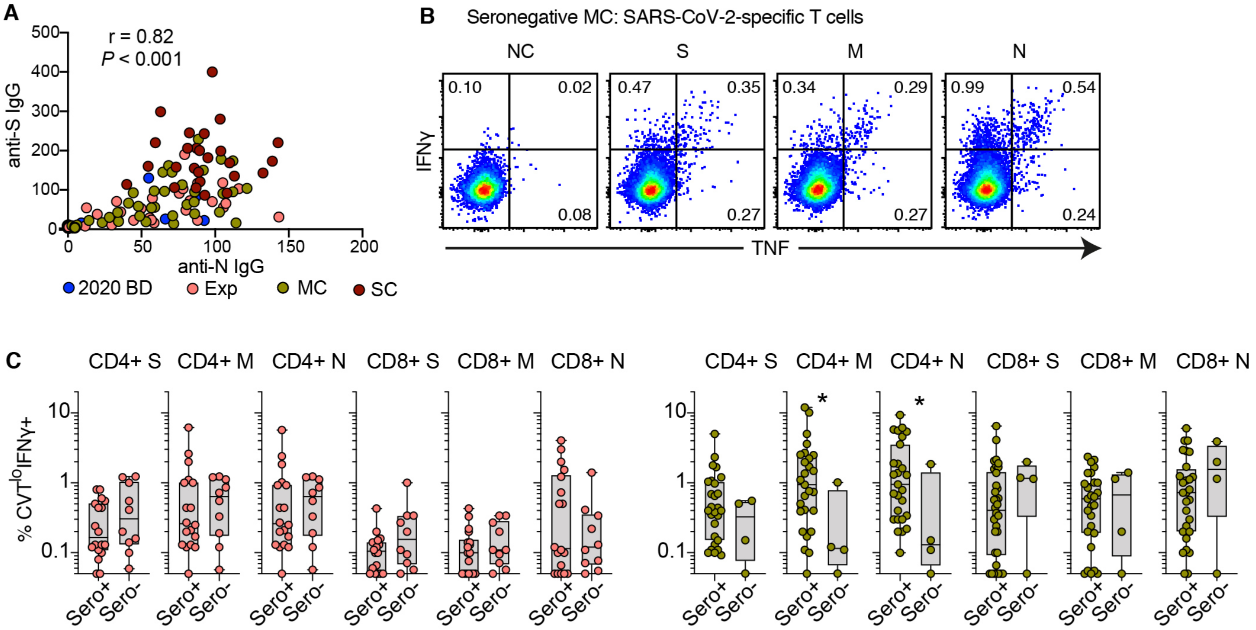


**Figure S8. Antibody correlations and SARS-CoV-2-specific memory T cell responses in seropositive and -negative convalescent COVID-19.** (**A**) Correlation between anti-nucleocapsid (N) and anti-spike (S) IgG levels. Each dot represents one donor. 2020 BD: Healthy blood donors from 2020. Exp: exposed family members. MC: individuals in the convalescent phase after asymptomatic/mild COVID-19. SC: individuals in the convalescent phase after severe COVID-19. (**B**) Representative flow cytometry plots showing functional SARS-CoV-2-specific memory T cell responses in a seronegative convalescent donor (group MC). Numbers indicate percentages in the drawn gates. NC: negative control. S: spike. M: membrane. N: nucleocapsid. **(C)** Comparative SARS-CoV-2-specific T cell analysis between seropositive and seronegative exposed family members (left) and asymptomatic/mild convalescence subjects. Related to Figure 4.
